# Glioblastoma stem cells express non-canonical proteins and exclusive mesenchymal-like or non-mesenchymal-like protein signatures

**DOI:** 10.1101/2022.02.06.479313

**Authors:** Haris Babačić, Silvia Galardi, Husen M. Umer, Deborah Cardinali, Serena Pellegatta, Mats Hellström, Lene Uhrbom, Nagaprathyusha Maturi, Alessandro Michienzi, Gianluca Trevisi, Annunziato Mangiola, Janne Lehtiö, Silvia Anna Ciafrè, Maria Pernemalm

## Abstract

Glioblastoma’s (GBM) origin, recurrence and resistance to treatment are driven by GBM cancer stem cells (GSCs). Existing transcriptomic characterisations of GBM classify the tumours to three subtypes: classical, proneural, and mesenchymal. The comprehension of how expression patterns of the GBM subtypes are reflected at global proteome level in GSCs is limited.

To characterise protein expression in GSCs, we performed in-depth proteogenomic analysis of patient-derived GSCs by RNA-sequencing and mass-spectrometry proteomics. We identified and quantified over 10,000 proteins in two independent GSCs panels, and propose a GSC-associated proteomic signature (GSAPS) that defines two distinct morphological conditions; one defined by a set of proteins expressed in non-mesenchymal - proneural and classical - GSCs (GPC-like), and another expressed in mesenchymal GSCs (GM-like). The expression of GM-like protein set in GBM tissue was associated with hypoxia, necrosis, recurrence, and worse overall survival in GBM patients.

In a proof-of-concept proteogenomic approach, we discovered 252 non-canonical peptides expressed in GSCs, i.e., protein sequences that are variant or derive from genome regions previously considered protein-non-coding. We report new variants of the heterogeneous ribonucleoproteins (HNRNPs), which are implicated in mRNA splicing. Furthermore, we show that per-gene mRNA-protein correlations in GSCs are moderate and vary compared to GBM tissue.

## Introduction

Glioblastoma (GBM) is the most common malignant primary brain tumour, inevitably fatal, and characterised by short survival after diagnosis with median overall survival (OS) at 15 months (Louis et al., 2016; Weller et al., 2015; Wen and Kesari, 2008). A widely accepted GBM classification, proposed by Verhaak *et al*. (2010), is based on mRNA expression patterns that distinguish four GBM subtypes: classical, mesenchymal, proneural, and neural (Roel G.W. Verhaak et al., 2010). More recently, the classification was revised by removing the neural subtype, and highlighting subtypes’ plasticity, i.e. the ability to switch from one to another (Wang et al., 2017). The CPTAC consortium has recently explored the protein expression in adult GBM tumours and proposed a multiomic classification of GBM tumour subtypes to nmf1 (proneural-like), nmf2 (mesenchymal-like), and nmf3 (classical-like) (Wang et al., 2021).

Extensive research about the origin of GBM has established the theory that cancer stem cells drive the development and progression of GBM, contribute to resistance to chemo- and radio-therapy, and induce GBM recurrence (Galli et al., 2004; Singh et al., 2003). Primary GBM stem cells (GSCs) have shown to reflect the diversity of GBM, recapitulate the tumour subtypes at mRNA level, and represent a good model to study the molecular profile of this cancer and explore new therapeutic targets (Johansson et al., 2020). Many efforts were undertaken to uncover gene expression signatures that are pivotal for GSC functions, expanding our understanding of the transcriptome and proteome of GBM and GSCs (Asif et al., 2019; Guardia et al., 2020; Johansson et al., 2020; Kozuka-Hata et al., 2012; MacLeod et al., 2019; Marziali et al., 2016; Mostovenko et al., 2018; Rheinbay et al., 2013; Song et al., 2017; Yanovich-Arad et al., 2021). Single-cell RNA-sequencing (scRNAseq) studies have demonstrated that GSCs are plastic and can switch between different subtypes^8^. Despite these efforts to characterize the transcriptional programs responsible for GSCs’ plasticity and stemness, no study has provided in-depth proteomic or proteogenomic profiling of primary GBM stem cells. Furthermore, it is not known how well GBM subtypes are recapitulated in GSCs at protein level.

The aim of this study was to explore the proteomic and proteogenomic landscape of GSCs, to enhance our comprehension on: (i) the molecular GSC phenotype at protein level; (ii) the relation between mRNA and protein levels in GSCs; (iii) whether GSC proteome expression is detectable at tissue level; and (iv) non-canonical peptides originating from genome regions previously considered as non-protein-coding.

Here, we report deep transcriptome and proteome profiling of patient-derived GSCs, by RNA-sequencing (RNAseq) and high-resolution isoelectric focusing coupled with liquid chromatography and mass-spectrometry (HiRIEF LC-MS/MS), respectively. We discovered a new GSC-associated protein signature (GSAPS), which we validated in an independent panel of GSCs from the HGCC cohort (Johansson et al., 2020), in primary and recurrent GBM tissue, and in another set of tumour tissue from the CPTAC GBM cohort (Wang et al., 2021) (Figure 1A). We demonstrate that the GSAPS recapitulates key features of GSCs, such as proneural-to-mesenchymal axis and hypoxic metabolism, consists of protein drug targets and has a potential association with OS in GBM. Furthermore, we report mRNA-protein correlations and non-canonical protein sequences expressed in GSCs, discovering potentially new protein-coding targets for research and treatment.

**Figure 1.**
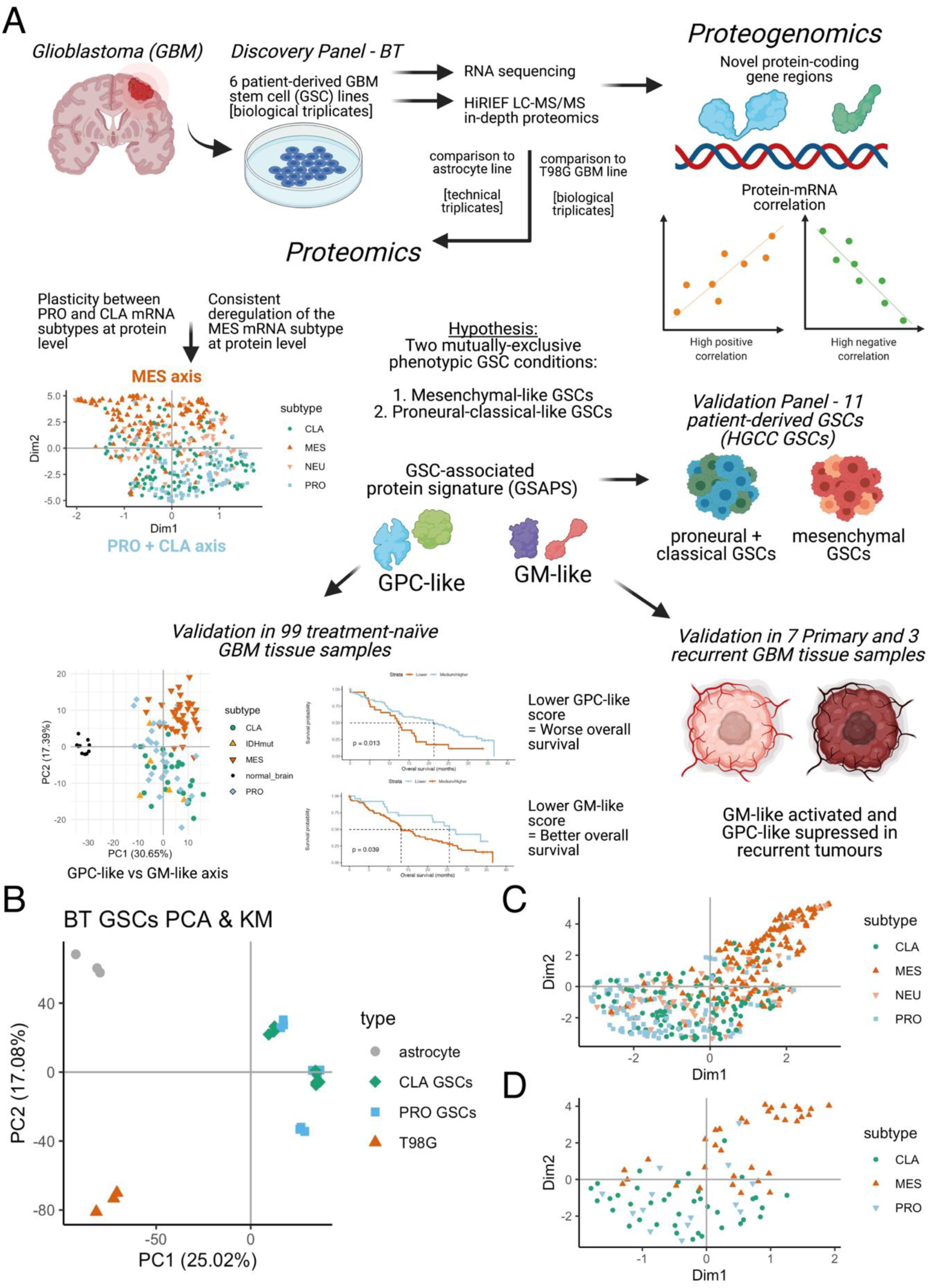
Study workflow and exploratory findings. **A.** In a discovery panel of six patient-derived GSC lines, previously subtyped as expressing the classical and proneural GBM subtype at mRNA level, we have identified variable enrichment of the proneural (PRO) and classical (CLA) GBM subtype, suggesting a plasticity between the two subtypes. However, all of the GSC lines had a suppression for the GBM mesenchymal (MES) subtype at protein level. We hypothesised that the GSCs are more distinctive at protein level based on whether they express the mesenchymal subtype or not and aimed to identify a protein signature (GSAPS), that consisted of two protein sets: the proneural+classical-like (GPC-like) protein set that was expressed in proneural and classical GSCs and a mesenchymal-like protein set (GM-like) expressed in mesenchymal GSCs. GSAPS was identifiable in another panel of 11 patient-derived GSCs, and in GBM tissue, where the expression of lower GPC-like protein scores was associated with worse overall survival, whereas lower GM-like protein scores were associated with better overall survival. Finally, by integrating proteomic and transcriptomic expression, we have performed proteogenomic analysis of the discovery panel of GSCs, discovering novel protein-coding gene regions and providing assessment of how well mRNA levels predict protein levels; **B.** PCA based on proteomic expression of the GSC samples and controls; **C-D.** UMAP of protein products of genes included in the Verhaak 2010 GBM subtypes’ gene sets (C) and Wang 2017 GBM subtypes’ gene sets (D).

## Results

### Protein identification and GBM proteome subtyping

To extract GSC-specific features, we selected six primary GSCs (hereafter referred to as BT GSCs) for RNAseq analysis and in-depth proteomic profiling (Figure 1A). Three GSCs were previously classified as expressing the classical subtype and three expressing the proneural subtype (Wang 2017 mRNA classification, Table S1) (De Bacco et al., 2021, 2012; Patanè et al., 2013). All samples were run in biological triplicates. For comparison, in the proteomic experiments we included primary human healthy astrocytes with three technical replicates representing normal brain cells, and three biological replicates of the T98G human glioblastoma cell line, representing a non-stem glioblastoma cell line (hereafter defined as controls). Across all samples, we identified 11,140 proteins, of which 9,161 proteins (82.24%) had no missing values and were included in the analyses. This is, so far, the most in-depth proteomic characterisation of GSCs.

Based on total proteome expression, the GSCs clearly clustered away from the normal brain cells as well as the T98G GBM cell line, and the GSC biological replicates clustered very close to each other (Figure 1B). Clustering the samples with proteins corresponding to genes previously implicated in GSC biology (Table S2) (Behnan et al., 2019; Codrici et al., 2016; De Bacco et al., 2012; Gimple et al., 2019; He et al., 2012; Lathia et al., 2015; Pointer et al., 2014; Prager et al., 2019; Saygin et al., 2019) showed clear separation of GSCs from the astrocyte and the T98G line (Figure S1A). Performing single-sample gene-set enrichment analysis (ssGSEA) to define the Verhaak GBM subtype at protein level showed that three cell lines overexpressed a different subtype at protein level compared to their initial mRNA subtype classification; some GSCs initially classified as proneural had enrichment for the classical subtype and vice versa. In addition, the proteins included in the proneural gene set projected closer to the classical gene set, suggesting that they were coexpressed in the GSCs (Figure 1C-D). This also implied that classical GSCs are more closely related to the proneural GSCs in human samples, as suggested in a mouse cell-of-origin gene signature in mouse GSCs (Jiang et al., 2017). The GSCs showed higher expression of protein products of genes included in both the proneural and classical subtype, but had a consistently lower expression of proteins deriving from mesenchymal gene sets, as compared to the non-stem cell lines (Figure S1B). Based on ssGSEA, we did not detect an activation of the Wang proneural and classical gene sets at protein level, possibly because these gene sets are smaller than the Verhaak GBM gene sets. However, all GSCs had a suppression for the Wang mesenchymal subtype, in agreement with the Verhaak gene sets (Figure S2A & S2B). Furthermore, the MET gene had consistent downregulation in BT GSCs, and all GSCs had higher EGFR to MET ratio compared to controls (Figure S2A & S2C), suggesting that higher EGFR-to-MET ratio and a downregulated MET could be biomarkers of the non-mesenchymal subtypes. The downregulation of MET in classical GSCs is in agreement with previous findings (De Bacco et al., 2012), however, we found that MET is downregulated in proneural GSCs at protein level opposing the findings of De Bacco et al. (2012).

### Protein-mRNA correlations in GSCs

Due to the variable protein expression of the gene sets used to classify GBM subtypes in the BT GSCs, we have continued with analysing the agreement between mRNA and protein per-gene products in GSCs. Per-gene correlation analysis of mRNA and protein matching to 9,007 genes showed an overall moderate agreement between mRNA and protein level (median Spearman’s *ρ* = 0.49, 5% FDR, Figure 2A, Table S3). Analysing several established GBM and splicing gene sets of interest (Beier et al., 2007; Uhlen et al., 2017; Roel G W Verhaak et al., 2010; Wang et al., 2017) showed similar mRNA-protein correlations as observed in the entire proteome identified in the GSCs (Figure 2B). Genes upregulated in glioma stem cells (Beier et al., 2007) and glioma-elevated genes (obtained from the Human Protein Atlas – HPA (Uhlen et al., 2017)) had a higher than overall mRNA-protein correlation, whereas genes involved in splicing and heterogeneous ribonucleoproteins (HNRNPs) had lower than overall correlation in GSCs.

**Figure 2.**
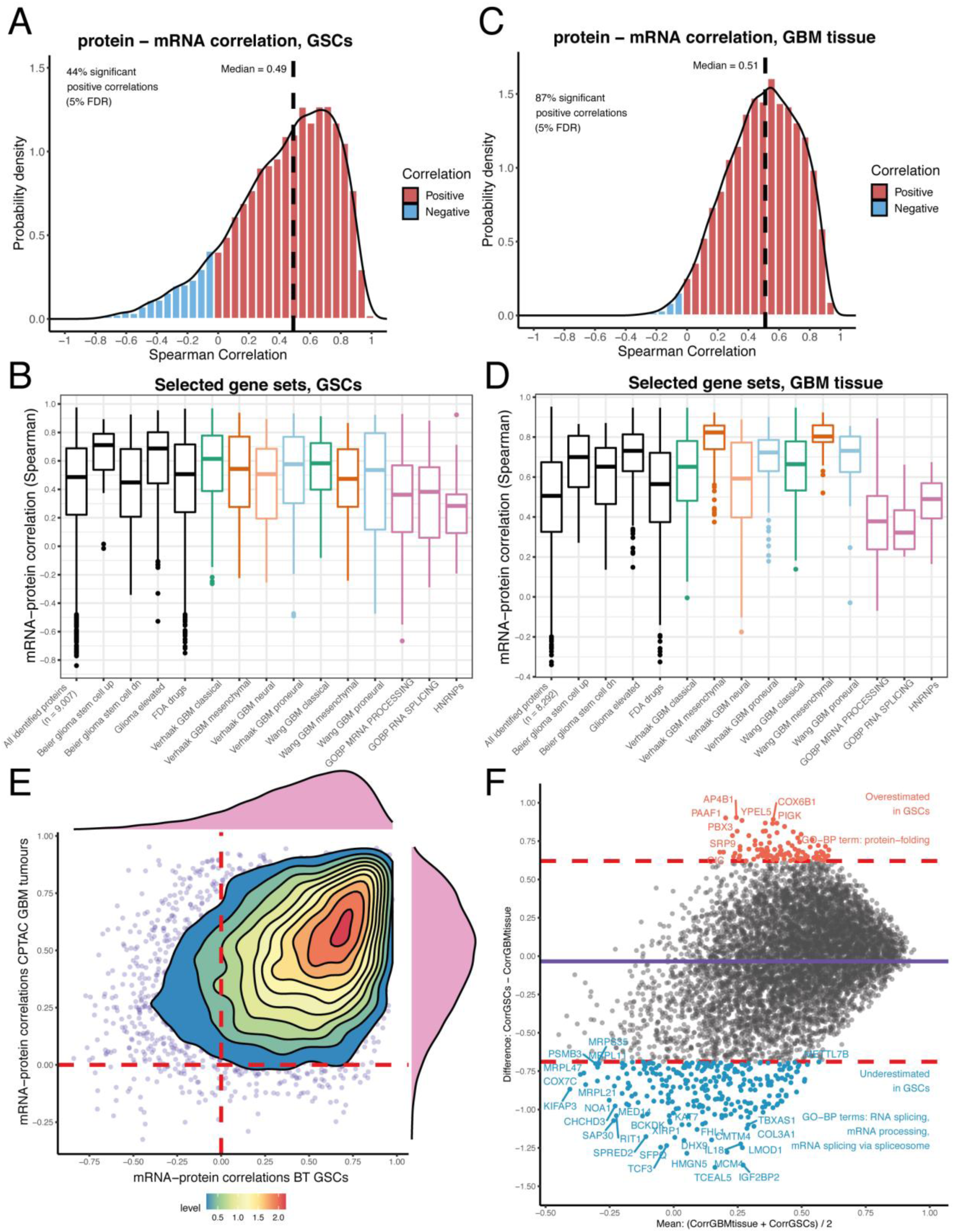
mRNA-protein correlations in BT GSCs and in CPTAC GBM tissue. **A.** mRNA-protein correlation of genes identified in BT GSCs with both RNAseq and HiRIEF LC-MS/MS; **B.** GSCs’ mRNA-protein correlation of genes included in selected gene sets of interest; **C.** mRNA-protein correlation of genes identified in GBM tissue with both RNAseq and HiRIEF LC-MS/MS, CPTAC cohort; **D.** GBM cancer tissue mRNA-protein correlation of genes included in selected gene sets of interest; **E.** Density plot comparing mRNA-protein correlation coefficients in GSCs and GBM tissue. Most of the genes have a positive correlation (> 0) in both GSCs and GBM tissue; **F.** Bland-Altman plot comparing the agreement between correlation coefficients in GSCs and in GBM tissue. The mean of the coefficients is plotted on the x axis and the difference between the coefficients is plotted on the y axis. The dashed lines show the 95% confidence intervals for the differences in correlation coefficients. Outside of the dashed lines are the genes with the largest disagreement in mRNA-protein correlations at GBM tissue and GSC level. The proteins below the lower dashed line having significantly lower mRNA-protein correlation in GSCs and proteins above the upper dashed line having significantly higher mRNA protein-correlation in GSCs, as compared to GBM tissue. These genes lists were enriched for the annotated gene ontology (GO) terms; the full enrichment terms are given in Figure S3.

In order to verify whether the overall moderate mRNA-protein correlations are observable at GBM tissue level as well, we downloaded proteomic and transcriptomic data from the recently published CPTAC GBM cohort, which includes proteomic profiling of 99 treatment-naïve GBM cancer tissues (Wang et al., 2021). Based on an analysis of mRNA and protein products deriving from 8,292 genes, GBM tissue also had a moderate overall mRNA-protein correlation (median Spearman’s *ρ* = 0.51, 5% FDR, Figure 2C, Table S4). GBM tissue had more statistically-significant correlations and less skewed distribution of mRNA-protein correlations compared to the GSCs, which is most likely due to the larger sample size of the GBM cohort that provided better estimates. The selected gene sets of interest showed mRNA-protein correlation patterns in tissue similar to those in GSCs (Figure 2D), and the majority of the genes had similar correlation between mRNA and protein in GSCs and GBM tissue (Figure 2E). Although most of the estimates in GSCs and GBM were in the same direction, a proportion of genes had varying mRNA-protein correlation in GSCs compared to GBM tissue. Comparing the agreement between correlations’ estimates by a Bland-Altman plot analysis showed that proteins involved in RNA splicing and protein-folding had lower and higher mRNA-protein correlation in the GSC lines compared to GBM tissue, respectively (Figure 2F, Figure S3), suggesting that GSCs have a higher degree of impaired splicing regulation but are less likely to accumulate unfolded proteins than GBM tissue due to better translation of proteins that regulate protein folding.

The mesenchymal gene sets had the highest concordance between mRNA and protein level in GBM tissue, with median correlation of r = 0.823 and r = 0.803 for the Verhaak and Wang mesenchymal gene sets, respectively (Figure 2D). This was much higher compared to the median mRNA-protein correlation in GSCs of r = 0.544 and r = 0.474 for the Verhaak and Wang mesenchymal gene sets, respectively (Figure 2B). The classical and proneural gene sets also had higher mRNA-protein correlation in GBM tissue, compared to GSCs, confirming that these gene sets might perform better at subtyping GBM tumours than subtyping GSCs. However, one limitation in our study is that the discovery panel did not include mesenchymal GSCs, which has possibly limited the variance in protein expression for the subtypes’ gene sets. Calculating the per-protein standard deviation in protein expression of genes included in the mesenchymal and classical gene sets showed higher variance in protein levels in GBM tissue than the GSCs, which could have driven the higher mRNA-protein correlations (Figure S4A & S4B). However, we found no such association for the proneural gene sets (Figure S4C), and the difference in the variance explained only 6-11% of the differences in the mRNA-protein correlations of the mesenchymal gene sets, which does not sufficiently explain the large overall difference between GBM tissue and GSCs in the mRNA-protein correlation for these gene sets. It is also likely that non-cancer cells, such as stromal and immune cells, could have contributed to a larger variance in protein expression of genes included in the GBM subtypes, suggesting that the GBM subtypes expression patterns might not be fully reflected at GSC level. The higher correlations in tissue for the mesenchymal gene sets are expected, because it has been recently shown that this subtype has a larger infiltration of immune cells (Wang et al., 2021). Still, other factors, such as gene sets’ size, mRNA decay, protein degradation, study sample size, protein identification and technical measurement errors could have all contributed to the disagreement in estimating mRNA and protein correlations in both GSCs and GBM tissue.

Overall, our findings demonstrate that the regulation of mRNA translation to protein follows similar patterns in GSCs as in GBM, and that GSCs can be a representative cell model for GBM to some degree. However, there was a notable disagreement between mRNA and protein levels, which warrants investigating the GSCs at the phenotypic level by analysing the proteome.

### GSC-associated protein signature reflects the proneural-mesenchymal axis

The plasticity between the classical and proneural subtype, flanked by the lack of consistent enrichment of the GBM gene sets at protein level, prompted us to hypothesise that gene programs associated with the proneural and classical subtype are coactivated in one type of GSCs. The suppression of the GBM mesenchymal subtype seemed a more consistent predictor for GSCs at protein level, leading to the hypothesis that GSCs at phenotypic (proteome) level express two exclusive programmes – either a mesenchymal-like or a proneural-classical-like.

To select a set of proteins that describes the hypothesised GSC phenotypes, we performed a differential expression analysis, to define a GSC-associated protein signature (GSAPS). We compared each GSC triplicate to each non-stem triplicate (astrocyte or T98G), to encompass the defining stem expression signature of each cell line, and selected the overlapping proteins that were consistently differentially expressed in the same direction in all BT GSCs and in all comparisons (Figure S5). This led to a core set of 524 proteins that we define as GSAPS (Figure 3A, Table S5).

**Figure 3.**
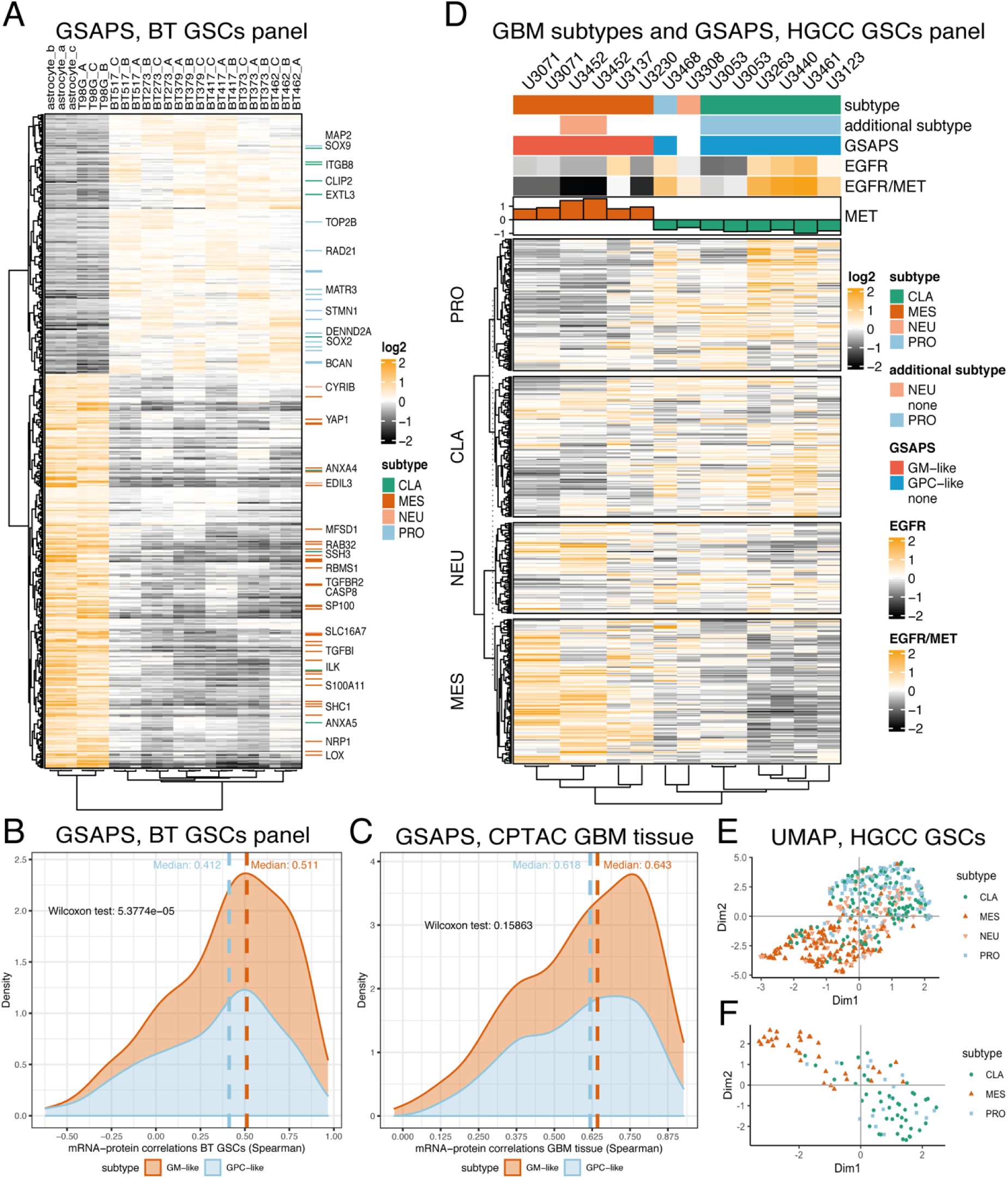
GSAPS and validation in the HGCC panel of GSCs. **A.** Hierarchical clustering of the BT GSC panel and controls with proteins included in GSAPS (distance: 1-Spearman’s r); **B.** Correlation between protein and mRNA levels of GSAPS protein sets in GSCs; **C.** Correlation between protein and mRNA levels of GSAPS protein sets in GBM tissue; **D.** Hierarchical clustering based on the protein expression levels of genes included in the Verhaak gene sets (distance: 1-Spearman’s r), and EGFR and MET protein expression in GSCs of different subtypes; **E-F.** UMAP dimensional reduction of the genes included in the Verhaak GBM subtypes’ gene sets (E) and the revised Wang GBM subtypes’ gene sets (F).

As expected, gene set enrichment analysis (GSEA) of GSAPS showed upregulation of the Verhaak proneural subtype gene set and downregulation of the mesenchymal subtype and the epithelial-to-mesenchymal transition (EMT) gene sets (Figure S6). Gliomas are not tumours derived from the epithelium, therefore the EMT is not directly applicable to them. However, a similar process, proneural-to-mesenchymal transition (PMT), has been described in GBM and is associated with worse prognosis and therapy resistance (Behnan et al., 2019; Bhat et al., 2013; Halliday et al., 2014; Phillips et al., 2006; Segerman et al., 2016; Wang et al., 2017). A predominant part of GSAPS consisted of upregulated proneural and downregulated mesenchymal markers, suggesting that the GSAPS is reflective of the PMT. The GSAPS also had the *hallmark* hypoxia gene set downregulated, along with several other hypoxia gene sets (Figure S6B), indicating that the BT GSCs were not hypoxic. The hypoxic metabolism has been associated with the mesenchymal subtype (Behnan et al., 2019; Tejero et al., 2019), suggesting that GSAPS can reflect the subtype-driven cellular metabolic condition.

Based on the differences in gene sets enriched among the upregulated and downregulated GSAPS proteins, we split it into two protein sets. The first consisted of the upregulated GSAPS proteins associated with the proneural signature, proliferation and non-hypoxic metabolism, henceforth referred to as GSAPS Proneural and Classical-like protein set (GPC-like), and the other consisted of the downregulated GSAPS proteins, associated with the mesenchymal signature, hypoxia, and EMT, henceforth referred to as GSAPS Mesenchymal-like protein set (GM-like). We hypothesised that these two protein sets define two different GSC conditions, which are mutually exclusive and would better define the specific stem phenotypes in GSCs than the previously established Verhaak and Wang gene signatures established for GBM tissue. Worth noticing is that 107 of the GSAPS proteins are targetable by FDA-approved drugs (31 in the GPC-like and 76 in GM-like set, Table S6), with some drugs targeting more than one protein in the signature and 33 drugs ongoing clinical trials in GBM (Table S7).

The overall protein-mRNA correlation of genes encoding for the GSAPS proteins was moderately positive (Spearman’s median r = 0.459), indicating that some features should be detectable at mRNA level but a large proportion of the GSC phenotype variance will be observable only at protein level. We detected a higher mRNA-protein agreement for genes included in the GM-like set in GSCs (Figure 3B), but this was not observed in GBM tissue (Figure 3C), which had higher mRNA-protein agreement for the GSAPS sets than GSCs.

### GSAPS defines two phenotypic conditions that differ in hypoxic metabolism

In order to confirm the GSAPS ability to define GSC conditions along the PMT axis, we performed proteomic expression profiling on another GSC panel consisting of GSCs of all GBM subtypes (Verhaak and Wang classification, based on mRNA expression). The extended cohort included 11 patient-derived GSC lines from the HGCC cohort, identifying 10,169 proteins across the cell lines, including cell lines classified as mesenchymal based on mRNA expression (Johansson et al., 2020). Subtyping the cell lines with single-sample GSEA (ssGSEA) at protein level showed that all GSCs that expressed the classical subtype also expressed the proneural subtype, and had a suppression for the mesenchymal subtype (Figure 3D), in line with previous observation from the BT GSCs. Clustering the proteins corresponding to the subtype-specific genes included in Verhaak (Roel G W Verhaak et al., 2010) and Wang (Wang et al., 2017) GBM gene sets showed again that the proteins included in the classical gene set projected closer to the proteins included in the proneural gene set and apart from the mesenchymal proteins (Figure 3E & 3F).

Applying GSAPS to the HGCC panel clustered the mesenchymal GSCs separately from proneural-classical GSCs (Figure S7). To further validate whether the GSAPS is reflective of PMT, we performed GSEA on the two GSAPS protein sets comparing the proneural-classical HGCC GSC lines to the mesenchymal GSC lines. As hypothesised, we detected a strong enrichment of both GSAPS protein sets (NES>3, p < 0.001), with the GPC-like set upregulated in the proneural-classical GSCs and the GM-like upregulated in the mesenchymal GSCs (Figure 4A and 4B). ssGSEA analyses showed that GSCs expressing the GPC-like phenotype had suppression of the GM-like phenotype, and vice versa, confirming the hypothesis that these conditions are mutually exclusive. GSEA comparing the protein expression of GPC-like GSCs to GM-like GSCs on *hallmark* gene sets showed metabolic differences between the cell lines, with GM-like GSCs enriched for hypoxia (Figure S8).

**Figure 4.**
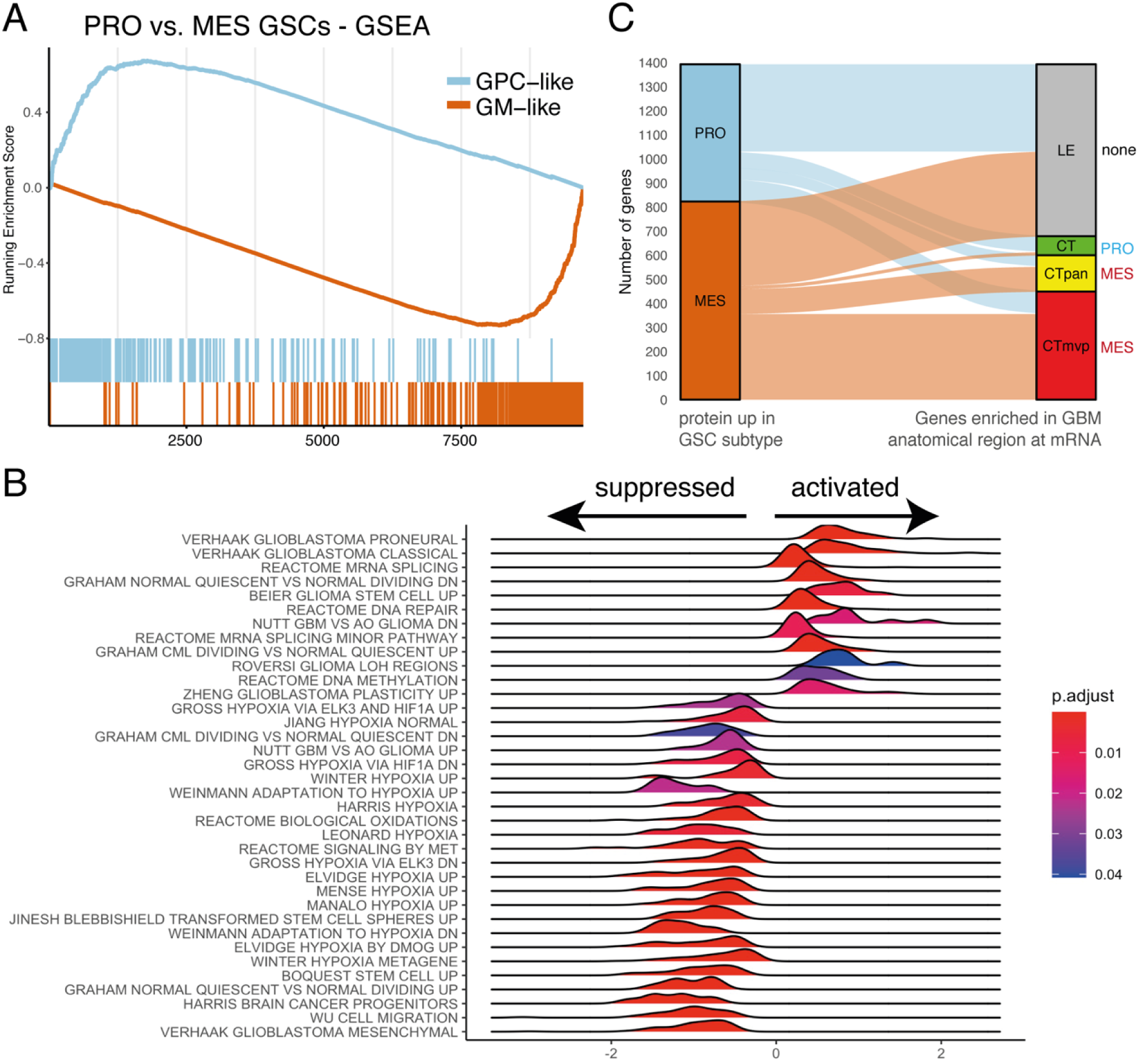
Proneural-mesenchymal axis and GSAPS association with different gene sets and pathways. **A.** GSEA of the GSAPS protein sets GPC-like and GM-like in proneural+classical GSCs as compared to mesenchymal GSCs. The GPC-like and GM-like protein sets were enriched in the proneural+classical GSCs and mesenchymal GSCs, respectively (NES > 3, p < 0.001, 1% FDR); **B.** MSigDb C2 gene sets (subcategory: GCP and REACTOME) enriched in the GPC-like GSCs as compared to GM-like GSCs at 5% FDR, GSEA, x axis = normalised enrichment score; **C.** Sankey diagram depicting the proportion of genes upregulated in the GPC-like (PRO) or GM-like (MES) GSCs at protein level that is enriched in different anatomical regions of GBM: leading edge (LE), cellular tumour (CT), cellular tumour palisading around necrosis (CTpan), and cellular tumour’s microvascular proliferation (CTmvp). On the right side of the diagram, the enriched protein set is annotated per region (two-sided Fisher’s exact test, p < 0.001, 1% FDR);

Considering that mesenchymal gene expression has been consistently associated with hypoxia, we hypothesised that the GM-like GSCs could be enriched in hypoxic regions of GBM tumour tissue, in proximity to necrosis, such as regions of tumour cells palisading around necrosis (CTpan) and tumour cells involved in microvascular proliferation (CTmvp). We then performed enrichment analysis comparing protein expression between GPC-like and GM-like GSCs to genes enriched in different GBM anatomical regions at mRNA level, based on the Ivy GBM Atlas (Puchalski et al., 2018) consisting of genes enriched at mRNA level in different GBM regions. Neither GSAPS set had enrichment in the leading edge (LE) region of GBM (OR = 1.058, Fisher’s test p = 0.485). However, genes overexpressed in the GM-like GSCs at protein level were enriched in regions of CTmvp (OR = 3.883, Fisher’s test < 2.2^-16^) and CTpan (OR = 2.115, Fisher’s test p = 1.19^-05^, Figure 4C). Oppositely, genes overexpressed in GPC-like GSCs at protein level were enriched in regions of cellular tumour – CT (OR = 5.259, Fisher’ test p = 4.149^-10^, Figure 4C). The findings suggest that GSCs adapt their phenotypic expression and thereby their subtype to local conditions, driving different elements of tumorigenesis and that the plasticity in itself is involved in driving the tumorigenesis. This is in line with previous observations within the Ivy GBM Atlas (Puchalski et al., 2018). However, one limitation is that the gene sets of the Ivy Atlas are derived by transcriptomic methods, leaving a gap to explore the regional protein expression in GBM for future endeavours.

In summary, these findings confirm that GSAPS is associated with PMT and that cultured GSCs exist in two mutually-exclusive phenotypic conditions, one characterised by the GPC-like protein set and another characterised by the GM-like protein set. The GSC phenotypes appeared enriched in different regions of the tumour.

### GSAPS is enriched in recurrent GBM tissue

Recurrent GBM tumours tend to have worse outcome and faster progression. Several studies have linked this to PMT, suggesting that proneural and classical GSCs are more sensitive to chemotherapy and radiotherapy, which eventually leads to selection and enrichment of the mesenchymal subtype within recurrent tumours (Behnan et al., 2019; Wang et al., 2017). To further demonstrate that GSAPS reflects PMT, we analysed 7 primary and 3 recurrent GBM tissue samples on proteomic level with HiRIEF LC-MS/MS, identifying 7,810 proteins, with 7,378 proteins quantified in all samples. One primary GBM tumour was excluded from analyses because it was highly necrotic on H&E staining and clustered separately from the other tumours (Figure S9A & S9B). GSEA between non-paired recurrent and primary tumours showed activation of the mesenchymal GBM gene set and suppression of pathways associated with GPC-like GSCs in recurrent tumours (Figure S9C & 9D). As hypothesised, GSEA on the GSAPS gene sets, comparing recurrent to primary GBM, showed a suppressed GPC-like and activated GM-like protein set in recurrent GBM tumours (Figure 5A). The GM-like protein set was also enriched in the necrotic GBM sample in ssGSEA (Figure S10), further indicating that the GM-like signature is associated with necrosis.

**Figure 5.**
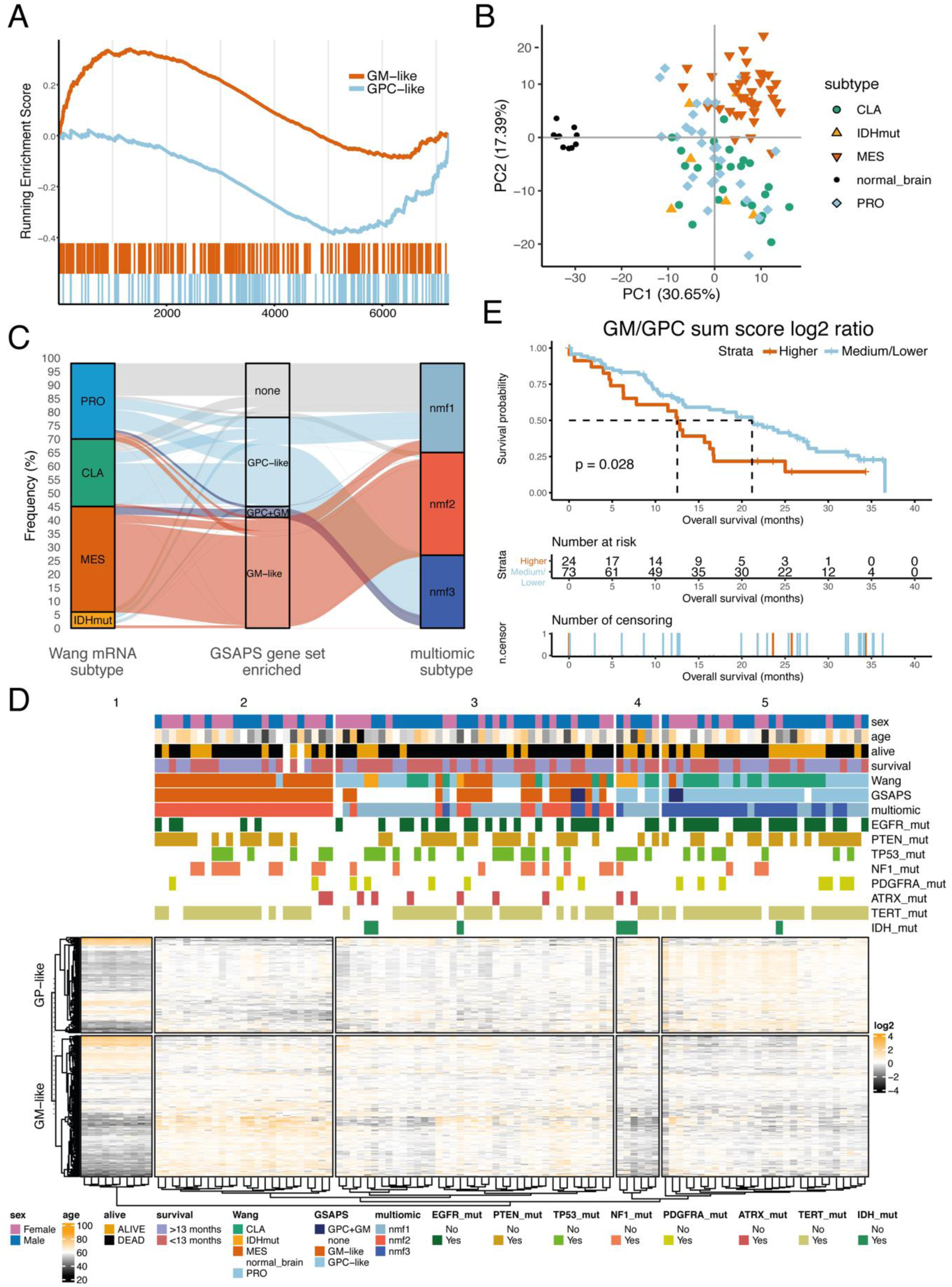
GSAPS expression in GBM tissue. **A.** GSEA on the GSAPS protein sets GPC-like and GM-like comparing recurrent to primary GBM tissue tumours (p < 0.001, 1% FDR); **B.** PCA clustering, based on log2 expression levels of proteins included in the GSAPS, of GBM tumours and normal brain tissue samples. GBM subtypes (mRNA, based on the Wang 2017 GBM classification(Wang et al., 2017)): CLA = classical, PRO = proneural; IDHmut = IDH-mutant tumour; MES = mesenchymal; **C.** Sankey diagram showing the proportion of GBM tumours of different transcriptomic subtypes (Wang 2017, GBM classification) that are enriched for the GSAPS protein sets GPC-like or GM-like or both, as compared to the CPTAC’s multiomic GBM subtypes recently described by Wang *et al*. (2021)(Wang et al., 2021); **D.** Hierarchical clustering of GBM tumours and normal brain samples based on the GSAPS (distance: 1-Spearman’s r). The different subtypes are shown in the annotation bars, as well as mutation status of common genomic markers in GBM; **E.** KM curves showing survival differences in patients categorised based on log2 GM to GPC protein sum score ratio to group of high (> third quartile) and medium/low (< third quartile) score ratios. The p-values are based on logrank tests; the dashed lines present the median overall survival in the corresponding groups.

### GSAPS protein signatures are associated with overall survival in GBM tissue

To demonstrate whether the GSAPS protein expression is maintained at tissue level, we explored its expression in GBM tumours from the CPTAC cohort. PCA analysis of 99 GBM tumours and 10 normal brain samples from the CPTAC cohort (Wang et al., 2021) based on GSAPS expression showed clear separation of GBM from normal brain and separated mesenchymal from non-mesenchymal cancer tissue (Figure 5B). The GPC-like protein set was enriched in GBM tumours of the classical or proneural subtype (Wang classification, mRNA (Wang et al., 2017)) and the multiomic nmf1 (proneural-like) or nmf3 (classical-like) subtype (Wang et al., 2021), whereas the GM-like protein set was enriched in GBM tumours of mesenchymal (Wang et al., 2017) and nmf2 (mesenchymal-like) multiomic subtype (Wang et al., 2021) (Figure 5C and 5D). This confirms that the previous observations in the CPTAC cohort are recapitulated in GSCs, i.e., mesenchymal GBM tumours exhibited a different proteomic and metabolomic profile from non-mesenchymal GBM tumours.

Considering that the GM-like signature was associated with hypoxia, necrosis and recurrence in GBM tissue, we hypothesised that it might be associated with worse OS in GBM. To prove the hypothesis, we calculated GPC and GM protein sum scores, by summarising relative expression of the proteins included in the corresponding GSAPS protein sets, and performed survival analysis. Adjusting for age, which was associated with worse OS in this GBM cohort, higher GPC protein sum scores had a statistically-non-significant association with longer OS (HR = 0.278, 95% CI: 0.067-1.15, likelihood ratio test (LRT), p=0.003, Table S8), whereas higher GM protein sum scores were associated with shorter OS (HR = 4.162, 95% CI: 1.181-14.662, LRT, p=0.001) in Cox proportional hazards models. This was also confirmed with KM survival analysis (Figure S11C & S11D, logrank test, p < 0.05). To incorporate both protein sets, we then calculated a log2 ratio of the GM to GPC protein sum score (log2 GM/GPC), which showed that higher log2 GM/GPC ratios were associated with worse OS (HR = 2.183, 95% CI: 1.063-4.481, LRT=0.002), adjusted for age in Cox models. This association remained consistent by categorising log2 GM/GPC ratio to higher (> third quartile) and medium/lower ratio (≤ third quartile) in KM curves (Figure 5E, logrank test, p = 0.028).

Overall, these results show that GSAPS describes a GSC cellular signal that can categorise tumours across the PMT axis, and that higher protein expression of the GM-like signature may be associated with worse OS in GBM.

### New protein-coding targets in GSCs

Stem cells often utilize parts of the genome that mature cells do not, such as early developmental genes, to obtain pluripotency. To explore if GSCs express *non-canonical proteins*, i.e. proteins expressed from genome regions considered as non-protein-coding, we employed a previously established proteogenomics pipeline (Umer et al., 2021; Zhu et al., 2018), to search for non-canonical peptides in BT GSCs. For this aim, we created an RNAseq-based database of predicted protein sequences, by translating the detected transcript sequences obtained from RNAseq of BT GSCs to protein sequences, and predicting corresponding tryptic peptides by *in silico* tryptic cleavage. We then appended the non-canonical database to a canonical database of protein sequences and searched for non-canonical peptides among the identified peptide spectra matches (PSMs). This approach allowed us to discover *novel non-canonical peptides* matching to novel protein sequences corresponding to genome regions predicted to be non-protein-coding, such as pseudogenes and lncRNAs, as well as *non-canonical peptides matching to canonical protein-coding genes* that have not been previously described, such as novel start sites, splice variants, gene extensions, etc.

We detected 252 non-canonical peptides expressed in the BT GSCs, half of them with 2 or more peptide spectral matches (>=2 PSMs, n = 118, 53.17%, Figure 6A, Table S9). More than half (53.97%) were novel peptides, whereas the remaining peptides matched to non-canonical sequences of protein-coding genes (Figure 6B & 6C).

**Figure 6.**
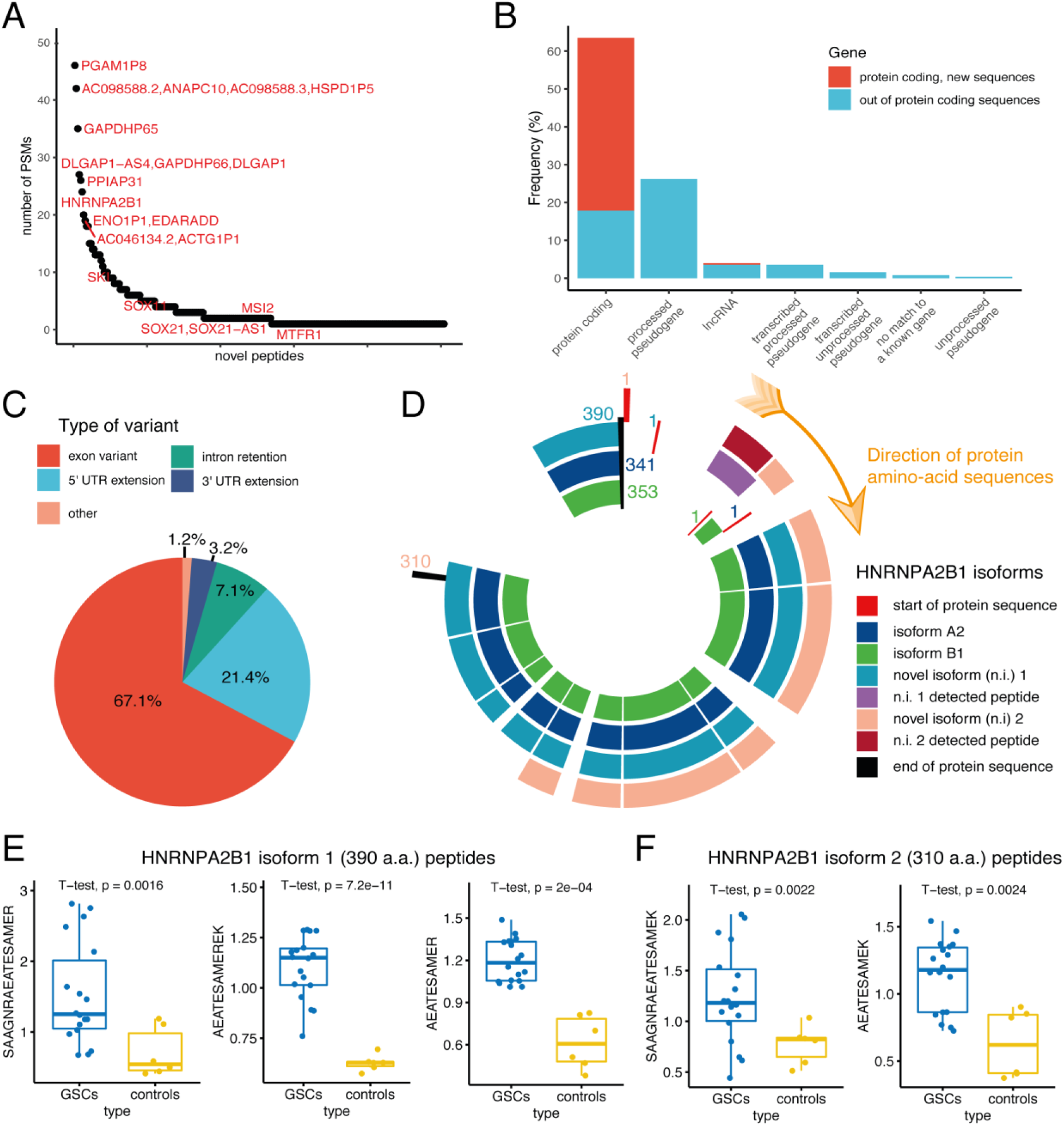
Non-canonical peptides expressed in GSCs. **A.** Number of PSMs per non-canonical peptide; **B.** Proportion of non-canonical and novel peptides classified according to matching gene type; **C.** Proportion of novel peptide classified according to the matching gene region; **D.** Canonical (A2 and B1) and non-canonical protein isoforms (here referred to as n.i. 1 and n.i. 2, both include 5’ extensions) of the HNRNPA2B1 gene. The plot shows the detected peptides positioned to the matching sequences of the canonical and novel isoforms of the HNRNPA2B1 gene. The numbers refer to the positions of the first and last amino acid of the corresponding isoform**; E.** Novel peptides matching to the novel protein isoform 1 of HNRNPA2B1 (390 amino acids long); **F.** Novel peptides matching to the novel protein isoform 2 of HNRNPA2B1 (310 amino acids long);

One tenth of the non-canonical peptides (n=23) matched to protein-coding genes included in the GSAPS, as expected mostly the GPC-like protein set (n=19), including exon variants of HNRNPA2B1, QKI, CUX1, EPHB3 and GAB1 (GPC-like) and 5’-UTR extensions of SOX2, TRIM24, QKI, and MSI2 (GPC-like), further highlighting their potential importance in GSC biology.

A recent screen of non-canonical open-reading frames characterised hundreds of new proteins in human induced pluripotent stem cells (iPSC) and human foreskin fibroblasts (HFF) (Chen et al., 2020). To validate the novel peptides discovered in our study, we downloaded the novel amino-acid sequences reported by Chen *et. al*, and found that 40 of the non-canonical peptides discovered in our study overlapped with their non-canonical protein sequences, providing independent support for these sequences (Table S9). Most of the non-canonical peptides were extensions of protein-coding genes (n = 33, 82.5%).

Sixteen non-canonical peptides found in GSCs matched to a family of ubiquitously expressed heterogenous nuclear ribonucleoproteins (HNRNP), which are involved in mRNA splicing, processing and metabolism^41^. Half of these peptides matched to processed pseudogenes (HNRNPA1-P8, -P12, - P14, -P16, and -P59), and the remaining half to variants of the isoforms A2 and B1. Among the non-canonical peptides matching to protein-coding genes, several matched to two novel protein-coding isoforms of HNRNPA2B1 reported by Chen *et al*. (2020), which have upstream extensions of the canonical protein isoforms’ sequences (Figure 6D) (Chen et al., 2020). The canonical HNRNPA2B1 protein had a higher expression in BT GSCs compared to non-stem controls and was part of the GPC-like protein set, along with other HNRNPs (HNRNP-U, -D, -DL, and -LL). Interestingly, the non-canonical peptides matching to the HNRNPA2B1 gene also had a higher expression in the BT GSCs compared to the controls (Figure 6E, Table S10, p < 0.05, 5% FDR), suggesting a role in GPC-like GSC biology. Still, it remains to be elucidated if the non-canonical protein sequences detected in GSCs in this study, such as those of HNRNPA2B1, are expressed at protein level only in GSCs or provide improved gene models. Overall, our findings show that some gene variants previously considered as non-coding are translated and expressed in GSCs and GBM at protein level and that a subset of these is related to proteins included in the GSAPS.

## Discussion

GBM is a highly malignant cancer, which is driven by GSCs and their ability to adapt in response to treatment and the tumour microenvironment. To improve treatment options for GBM patients, it is essential to understand the underlying mechanisms driving GSCs and how mRNA is translated to protein level, allowing the tumour to progress, adapt, and resist therapeutic interventions.

In this study we have performed the most in-depth proteogenomic analysis of GSCs to date, providing a new layer of information on GSC biology. Based on HiRIEF LC-MS/MS proteomics of primary GSCs we present a new GSC-associated protein signature (GSAPS). GSAPS recapitulates GSC-specific features, such as PMT and hypoxia, and was validated in an independent panel of GSCs from the HGCC cohort (Johansson et al., 2020). We discovered that non-mesenchymal GSCs express proteins belonging to both the proneural and classical subtype, maintaining the expression of a core set of proteins that we defined as GPC-like. On the other side of the spectrum were the mesenchymal-like GSCs, enriched for the GM-like protein set, with MET among proteins with highest levels. In line with the PMT hypothesis, we find that GSCs mainly exist in two phenotypic conditions, one defined by the GPC-like signature and another defined by the GM-like signature, which are mutually exclusive.

Furthermore, we show that the GM-like set is enriched in recurrent GBM tissue, in regions characterised by hypoxia and necrosis, and mesenchymal GBM tumours. Previous observations at tissue proteome level from the CPTAC cohort (Wang et al., 2021), have shown that mesenchymal GBM tumours have higher MET levels and are enriched for EMT, hypoxia, glycolysis, angiogenesis and inflammatory pathways, however it was not clear whether the pathways were enriched in GBM tumour cells or due to microenvironment. We demonstrated that all these observations at tissue level are driven by expression patterns at GBM cellular level, adding to our understanding of GBM tissue development.

From a clinical perspective, the GSAPS encompasses over 100 protein drug targets, out of which 33 are currently undergoing clinical trials for GBM. It is tempting to speculate that the signature might serve a purpose in drug-development guidance, where a combination treatment targeting proteins in both the GPC-like and the GM-like protein set might be more effective. Furthermore, we report that higher GM over GPC ratio might be associated with worse OS in GBM and could be of prognostic value. However, this has to be confirmed in larger, independent cohorts.

Possible limitations of our work are in the initial derivation of GSAPS by comparing proneural and classical GSCs to two types of non-stem cell lines instead of using several non-stem controls, and that the cells are cultivated *in vitro*. However, we have stringently filtered the proteins that would define GSAPS, and the strength of the *in vitro* cultivation of GSCs is in the ability to use them to experimentally validate their stem-cell nature through established methods described elsewhere (De Bacco et al., 2021). It is also relevant to point out that GSAPS showed consistent findings in subsequent validation of the signature in another panel of validated GSCs that use different culturing and stem-cell validation methods (Johansson et al., 2020). Furthermore, we have shown that the GSAPS expression can be traced in GBM tissue in our experiments and in a publicly available dataset on GBM tissue proteome expression, potentially providing clinically-useful information.

In summary, we present the most in-depth proteogenomic characterisation of GSCs to date, and report a new GSC-associated protein signature that differentiates two phenotypic conditions of GSCs along the proneural-to-mesenchymal axis. We have shown that some phenotypic patterns enriched in GBM subtypes at tissue proteome level are driven by protein expression programmes at GBM cellular level. Finally, we discover novel protein-coding gene regions in GSCs, some of which have been reported in other, non-cancer cells and some that are uniquely reported in this study. These findings allow studying GBM at a GSC cellular proteomic level, improve our understanding of GSC biology and identify new, both protein and pathway-related, subtype-specific therapeutic targets for GSCs.

### Key points

- This study provides the most in-depth proteome analysis of GSCs to date, comparing protein to mRNA levels. Only a subset of proteins has high correlation to mRNA levels.
- Two protein sets define a GSC-associated protein signature that distinguishes two phenotypic conditions of GSCs, which are mutually exclusive and have an inverse association with clinical outcomes in GBM.
- In GSCs, we discovered protein sequences matching to genes previously established as non-protein-coding. These novel non-canonical proteins, along with newly discovered variants of protein-coding genes in this study, may have implications in GBM.

### Importance of the study

By identifying over 10,000 proteins in two patient-derived GSC panels, this study is the most in-depth data resource of protein expression in GSCs, to date. Overall, mRNA levels are moderately good at predicting protein levels, highlighting the importance of understanding protein expression signatures behind GSC phenotypes.

We report a new GSC-associated protein signature (GSAPS) that describes two phenotypic conditions of GSCs. The expression at tissue level of the two protein sets that consist GSAPS, i.e., the GPC-like and GM-like sets, had an inverse association with clinically-relevant outcomes in GBM, such as necrosis, recurrence and overall survival, and may identify new treatment targets.

Proteogenomics allows for discovering non-canonical protein sequences that have not been observed before, matching to new protein variants, or pseudogenes and long-non-coding RNAs not expected to be protein-coding. The discovery of non-canonical proteins in GSCs questions established gene models and indicates potentially new proteins, which may have implications in GBM and warrant further investigation.

## Materials and methods

### GSCs and GBM tumours

BT human GSC lines were grown as neurospheres in serum-free medium as described and validated in (De Bacco et al., 2021; Galli et al., 2004). The human glioblastoma T98G cell line and the astrocyte line were purchased from ATCC and CliniSciences (Guidonia Montecelio, Italy), respectively. The HGCC human GSC cultures are part of the HGCC biobank (https://hgcc.se/), and have been previously described and validated (Johansson et al., 2020; Xie et al., 2015a).

Ten GBM tissue samples were collected and fresh-frozen at the Neurosurgical Unit and evaluated by a pathologist at the Hospital Spirito Santo, Pescara, “G. D’Annunzio” University, Chieti, Italy. All patients gave informed consent and the samples were collected and processed in accordance with the Declaration of Helsinki. The molecular analyses were approved by the Ethics Committee of the Provinces of Chieti and Pescara and of the “G. d’Annunzio” University of Chieti-Pescara.

### Cell culture

The T98G cells were cultured in DMEM supplemented with 10% foetal bovine serum. The astrocytes were cultured in Astrocyte Medium (ScienCell #1801). We grew three biological replicates for each BT GSC and the T98G line, and one biological replicate for the astrocyte line.

Handling of HGCC human tissues and data were performed in accordance with the protocol approved by Uppsala ethical review board (2007/353) and informed written consent was obtained from all patients. The cells were cultured as previously described (Jiang et al., 2017; Xie et al., 2015b), and were analysed between passage 10-19. Briefly, cultures were maintained on poly-ornithine/laminin-coated dishes in DMEM/F12 Glutamax (Gibco) and Neurobasal medium (Gibco) mixed 1:1 with addition of 1% B27 (Invitrogen), 0.5% N2 (Invitrogen), 1% Penicillin/Streptomycin (Sigma), 10 ng/ml each of EGF and FGF2 (Peprotech). They have been regularly screened for mycoplasma infection using a PCR-based method with the primers Myco1 (5’-GGCGAATGGGTGAGTAACACG) and Myco2 (5’-CGGATAACGCTTGCGACTATG) (Invitrogen) and no cultures have tested positive.

### GBM tumour tissue processing

The tissue samples were fixated on OCT and cut into 10 μm-thick sections, of which 30 sections were collected in a tube for lysis and parallel sections were fixed on slides for haematoxylin and eosin staining. The sections collected in tubes were washed in PBS to remove the blood, centrifuged, and the tissue pellets were used for subsequent DNA, RNA, and protein isolation with the AllPrep DNA/RNA/Protein Mini Kit (Qiagen).

### RNA sequencing

Sequencing libraries for whole transcriptome analysis were prepared using Stranded mRNA-Seq Library Preparation Kit. RNA-seq was performed on an Illumina HiSeq 2500 Sequencer using standard conditions at the Next Generation Sequence Facility of University of Trento (CIBIO).

#### RNA isolation, library preparation, RNA-sequencing, qRT–PCR

Total RNA from the BT GSCs was isolated by TRIzol (Invitrogen), subjected to DNase-I (Ambion) treatment and RNAs were depleted of ribosomal RNA. Two RNA samples derived from normal brain were purchased from Clontech Laboratories and BioChain respectively.

#### Data Quality check

The fastq files generated by the Illumina sequencer were monitored for quality by the FastQC tool (https://www.binformatics.babraham.ac.uk/projects/fastqc/, version 0.11.6). It provides a modular set of analyses which tests if the data has any problems. Since for each sample there is one FASTQC output, with several results, it was decided to use multiQC tool (http://multiqc.info/, v.1.4) to aggregate the information for a better interpretation. The main outcome of these analysis is that the reads have very good quality and despite some differences among samples, the further analyses could be done without corrections at this stage.

#### Transcript quantification

Transcript quantification was performed using Salmon (Patro et al., 2017). Salmon applies a quasi-mapping with a two-phase inference procedure to quantify expression at the transcript level. The unique feature that distinct Salmon from other transcript assemblers account is in its ability to account for experimental and other biases that are common to RNA-seq data such GC content. ENSEMBL cDNA release 99 from GRCh38 was used as the target transcriptome. To obtain gene-level quantifications, the median value across the transcripts of each gene was assigned as the gene expression. All options were set to default and -l A parameter was set to detect the library type from the RNA-seq datasets.

### Mass-spectrometry-based proteomics

The samples were prepared and run following the HiRIEF LC-MS/MS protocol, as previously described (Branca et al., 2014).

#### Cell lysis and in-solution digestion

The BT GSCs cells were lysed in 200 μl SDS-lysis buffer (containing (4% (w/v) SDS, 50 mM HEPES pH 7.6, and 1mM dithiothreitol) using 1:4-10 of sample to buffer ratio. Afterwards, the cells were heated at 95°C for 5 min while shaking on a pre-warmed block, and sonicated to dissolve the pellet and disrupt the remaining DNA. The lysate was then centrifugated at 14 000xg for 15 min and the supernatant removed. Proteins from HGCC cells and GBM tissue were extracted with the AllPrep DNA/RNA/Protein Mini Kit (Qiagen).

The protein concentration in the lysate was determined by Bio-Rad DC Assay and equal amounts of each sample was subjected to in-solution digestion. Briefly, the cell pellet was denatured at 95°C for 5 minutes followed by reduction with dithiothreitol and alkylation with chloroacetamide at end concentrations of 5mM and 10mM respectively. LysC was added at a 1:50 (w/w) ratio and digestion was performed at 37°C 6 hours or overnight. The samples were further digested by trypsin at a 1:50 (w/w) ratio with 37°C overnight incubation. After LysC/trypsin digestion, ∼1% of each peptide sample was aliquoted for ∼15min gradient LC-MS/MS runs to check for protease activity by the samples’ miscleavage rate.

#### TMT-labelling

Before labelling, equal amounts of peptide samples were pH adjusted using TEAB, pH 8.5. The resulting peptide mixtures were labelled with isobaric TMT-tags (Thermo Scientific). Biological triplicates of the BT GSCs and the T98G line, and technical triplicates of the astrocyte line were labelled with three TMT-10-plex sets, using two internal standards per set. The internal standards were made of sample pools. HGCC GSC samples were run in one TMTpro-16-plex set, without an internal standard, leaving the 133C and 134N channels empty. GBM tissue samples were labelled with one TMT-10 set, without an internal standard. Labelling efficiency was determined by LC-MS/MS before pooling of samples. Subsequently, sample clean-up was performed by solid phase extraction (SPE strata-X-C, Phenomenex). The labelling schemes per sets can be found in Tables SM1A, SM1B, and SM1C (Supplementary File 3).

#### High resolution isoelectric focusing (HiRIEF)

The HiRIEF prefractionation method at peptide level was applied as previous described (Branca et al., 2014). Briefly, after sample clean-up by solid phase extraction (SPE strata-X-C, Phenomenex), the sample pool was subjected to peptide IEF-IPG (isoelectric focusing by immobilized pH gradient) in pI range 3-10 (1mg). For the proteogenomics experiments, the sample pools of the BT cells were subjected to additional IEF and LC-MS/MS run in a separate experiment on IPG strips in the pI range 3.7-4.9, to increase the detection of peptides. The freeze-dried peptide sample was dissolved in 250µl rehydration solution containing 8M urea, and allowed to adsorb to the gel strip by swelling overnight. The 24cm linear gradient IPG strip (GE Healthcare) was incubated overnight in 8M rehydration solution containing 1% IPG pharmalyte pH3-10 (GE Healthcare). After focusing, the peptides were passively eluted into 72 contiguous fractions with MilliQ water / 35% acetonitrile / 35% acetonitrile and 0.1% formic acid, using an in-house constructed IPG extractor robotics (GE Healthcare Biosciences AB, prototype instrument) into a 96-well plate (V-bottom, Greiner product #651201). The BT GSCs samples were rerun and additionally fractionated by IEF-IPG in pI range 3.7-4.9, in order to detect more peptides for proteogenomic analyses. The resulting fractions were then freeze dried and kept at -20°C until LC-MS/MS analysis.

#### LC-MS/MS analysis

Online LC-MS was performed using a Dionex UltiMate™ 3000 RSLCnano System coupled to a Q-Exactive HF mass spectrometer (Thermo Scientific). Each plate well was dissolved in 20 ul solvent A and 10 ul were injected. Samples were trapped on a C18 guard-desalting column (Acclaim PepMap 100, 75μm x 2 cm, nanoViper, C18, 5 µm, 100Å), and separated on a 50cm long C18 column (Easy spray PepMap RSLC, C18, 2 μm, 100Å, 75 μm x 50 cm). The nano capillary solvent A was 95% water, 5% DMSO, 0.1% formic acid; and solvent B was 5% water, 5% DMSO, 95% acetonitrile, 0.1% formic acid. At a constant flow of 0.25 μl min^−1^, the curved gradient went from 2% B up to 40% B in each fraction, followed by a steep increase to 100% B in 5 min and subsequent re-equilibration with 2% B.

FTMS master scans with 60,000 resolution (and mass range 300-1700 m/z) were followed by data-dependent MS/MS (30 000 resolution) on the top 5 ions using higher energy collision dissociation (HCD) at 30% normalized collision energy. Precursors were isolated with a 2 m/z window. Automatic gain control (AGC) targets were 1e6 for MS1 and 1e5 for MS2, with minimum AGC target of 1e3. Maximum injection times were 100 ms for MS1 and 100 ms for MS2. The entire duty cycle lasted ∼2.5 s. Dynamic exclusion was used with 30.0s duration. Precursors with unassigned charge state or charge state 1, 7, 8, >8 were excluded.

#### Protein identification

Raw MS/MS files were converted to mzML format using msconvert from the ProteoWizard tool suite(Kessner et al., 2008). Spectra were then searched in the Galaxy framework using tools from the Galaxy-P project (Boekel et al., 2015; Goecks et al., 2010), including MSGF+ (Kim and Pevzner, 2014) (v2020.03.14) and Percolator (Kall et al., 2007) (v3.04.0), where 8 subsequent HiRIEF search result fractions were grouped for Percolator target/decoy analysis. Peptide and PSM FDR were recalculated after merging the percolator groups of 8 search results into one result per TMT set. The reference database used was the human protein subset of ENSEMBL101. Quantification of isobaric reporter ions was done using OpenMS project’s IsobaricAnalyzer (Rost et al., 2016) (v2.5.0). Quantification on reporter ions in MS2 was for both protein and peptide level quantification based on median of PSM ratios, limited to PSMs mapping only to one protein and with an FDR q-value < 0.01. FDR for protein level identities was calculated using the -log10 of best-peptide q-value as a score. The search settings included enzymatic cleavage of proteins to peptides using trypsin limited to fully tryptic peptides. Carbamidomethylation of cysteine was specified as a fixed modification. The minimum peptide length was specified to be 6 amino acids. Variable modification was oxidation of methionine.

### Proteogenomic identification

The proteogenomic pipeline is described elsewhere in detail, a brief description is provided as follows(Umer et al., 2021). Transcripts were assembled from the RNA-seq data of each sample using stringTie (version 2.113) (Kovaka et al., 2019) based on the human reference gene annotations (ENSEMBL99). Next, transcripts with low expression level (TPM <1) were removed and a peptide database was generated from the transcript sequences using custom scripts. Tryptic peptides with a minimum length of eight amino acids and a maximum length of 40 amino acids were kept. The database was fractionated based on the peptide isoelectric points as further detailed in (Branca et al., 2014). Finally, the human canonical proteins (ENSEMBL99) were appended to the peptide database.

The proteomics data from each cohort were searched against the peptide database from the same cohort using MS-GF+ Release (version 15 January 2020). Percolator (version 3.04.0) was used for Percolator target-decoy scoring. Peptides at FDR<1% were considered significant, while those matching canonical protein sequences were removed. Using BLAST, the remaining peptides were searched against a larger collection of reference protein databases that included Uniprot version 11 December 2019, Gencode version 33, Ensembl version 99, and RefSeq (version 29 May 2020). Peptides matching any sequence were removed and those with one mismatch were further validated using SpectrumAI (Zhu et al., 2018). Finally, the list of novel peptides contained peptides with more than one mismatch or no match to known proteins as well as those that passed SpectrumAI.

### Bioinformatics and statistical analyses

#### Differential expression and GSAPS algorithm

Protein or peptide differential expression was performed with a two-sided t test for all comparisons and corrected for multiple testing with the false discovery rate (FDR), at 5%. The GSAPS was isolated by comparing each BT GSCs triplicate to a control (astrocyte or T98G line) triplicate, and finding the intersect of proteins consistently upregulated and downregulated in the BT GSCs as compared to controls (see Figure S5).

#### Protein-mRNA correlation

Protein per-gene expression was calculated as the average of the proteins matching to the same gene, whereas the mRNA per-gene expression was calculated as the sum of TPMs per gene. Correlations between matching protein and mRNA expression levels per overlapping genes were tested with the Spearman’s correlation coefficient and permutation test at alpha = 0.05, and corrected for multiple testing with the FDR. Protein-mRNA correlation for the CPTAC data was performed using processed and normalised proteomic and transcriptomic data available from (Wang et al., 2021). The selected gene sets were extracted from the MSigDb database (Liberzon et al., 2015; Subramanian et al., 2005), apart from the ‘Glioma-elevated’ and ‘FDA drugs’ datasets, which were extracted from the Human Protein Atlas (Uhlen et al., 2017).

The Bland-Altman analysis on agreement in correlations between GSCs and GBM tissue was performed as previously described (Bland and Altman, 1986). The genes outside the 95% CI of the Bland-Alman plot were considered to have strong disagreement; we extracted the gene lists above and below the 95% CI and performed enrichment analysis with an overrepresentation test in g:Profiler.

#### Feature reduction, visual projection and clustering

PCA, UMAP, and hierarchical clustering of samples based on protein expression was performed on scaled log2 relative protein expression values. We used the prcomp, umap, and Heatmap functions from the stats, umap, and ComplexHeatmap packages, respectively.

#### ssGSEA, GSEA and MSigDB

ssGSEA was performed by ordering the protein rank according to their log2 relative protein expression values in a sample and performing a GSEA on gene sets of interest, adjusting for multiple comparisons at 5% FDR. For subtyping the GSCs, the Verhaak gene sets were downloaded from the MSigDB database (Liberzon et al., 2015; Subramanian et al., 2005) and we created a dataset with Entrez IDs for the Wang gene sets and the GSAPS protein sets. GSEA analyses were performed separately for published, hallmark, and GO biological processes’ gene sets by sub-setting the MSigDB to the C2 GCP and REACTOME, H, and C5 Biological processes categories. The ranking in the comparisons GPC-like vs. GM-like GSCs and recurrent vs. primary GBM tissue was based on the difference in log2 average expression in the first group and the log2 average expression in the second group. For all the GSE analyses we used the GSEA function from the clusterProfiler package.

#### *In silico* validation

##### GBM anatomical localisation

GBM differentially expressed gene sets per anatomic region were downloaded from the Ivy League GBM Atlas (Puchalski et al., 2018), including gene sets of leading edge (n=1,998), cellular tumour (n=114), palisades around necrosis (n=389), and microvascular proliferation (n=1,126). The gene sets per regions consisted of genes two-folds (log2-FC > 1) differentially expressed in that region as compared to the remaining regions, at 1% FDR, based on an edgeR analysis. We calculated the mean protein log2-FC between GPC-like and GM-like HGCC GSCs as a difference between mean log2 protein values and categorised them as up in GPC-like (if log2-FC>0) and up in GM-like (if log2-FC<0). We then made contingency tables and tested if the proteins were overrepresented in the anatomical regions’ gene sets with a two-sided Fisher’s exact test, at alpha < 0.05 and at 5% FDR.

##### CPTAC proteomics dataset

Processed, mass-spectrometry global-proteomics, log2-normalised protein expression data of GBM tissue samples (n = 99) and normal brain tissue samples (n = 10), along with clinical, subtype, molecular and survival data were downloaded from the CPTAC cohort (Wang et al., 2021). Based on the expression of proteins included in the GSAPS, the samples were clustered with PCA and hierarchical clustering (method: Euclidean distance). We then performed ssGSEA for the GPC-like and GM-like protein set, by ranking the proteins within a sample based on their log2 relative expression values.

We first performed survival analyses with Kaplan-Meier (KM) curves and a log-rank test at alpha=0.05, categorising the GBM patients according to GSAPS gene set enrichment (at protein level). The overall survival was calculated as the time period from date of initial pathological diagnosis to date of death or date of loss to follow-up. GPC and GM sum scores were calculated by summing up the relative protein expression values of the proteins included in the GPC-like and GM-like protein set, respectively, and log2-normalising them. We then performed a survival analysis with Cox proportional hazards models and a likelihood ratio test at alpha = 0.05, adjusting the scores for age. To further confirm the association between the GPC and GM sum scores, we categorised them based on quartile expression values to high/medium (> first quartile) and low score (< first quartile) and performed KM survival analysis with a logrank test, at alpha = 0.05. Finally, we calculated a log2 ratio of the GM to the GPC sum score and performed survival analysis both with Cox proportional hazards models and likelihood ratio test, adjusting for age. We then categorised the GM/GPC ratio to high (> third quartile) and low/medium (< third quartile) and performed KM survival analysis with a logrank test, at alpha = 0.05.

##### Software

All analyses were performed in R v.4.0.3.

## Data availability

The mass spectrometry proteomics data have been deposited to the ProteomeXchange Consortium via the PRIDE partner repository with the dataset identifiers: PXD027341, PXD027339 and PXD027335. RNAseq files, the datasets and the code used for the analyses can be provided by the corresponding authors upon reasonable request.

## Supplementary files

Supplementary File 1 – Supplementary figures: Figure S1-S11.

Supplementary File 2 – Supplementary tables: Table S1-S10.

Supplementary File 3 – Supplementary methods tables: Table SM1A-1C.

## Acknowledgments

We acknowledge support from the Proteogenomics Facility at Science for Life Laboratory in Stockholm, Sweden, with special gratitude to Dr Xiaofang Cao and Dr Georgios Mermelekas. We thank Dr Gianluca Sala from University of Chieti-Pescara for his help in providing the GBM tissue samples, and to Dr. Fabio Socciarelli for assistance with H&E staining.

## Supplementary Figures to manuscript

**Figure S1.**
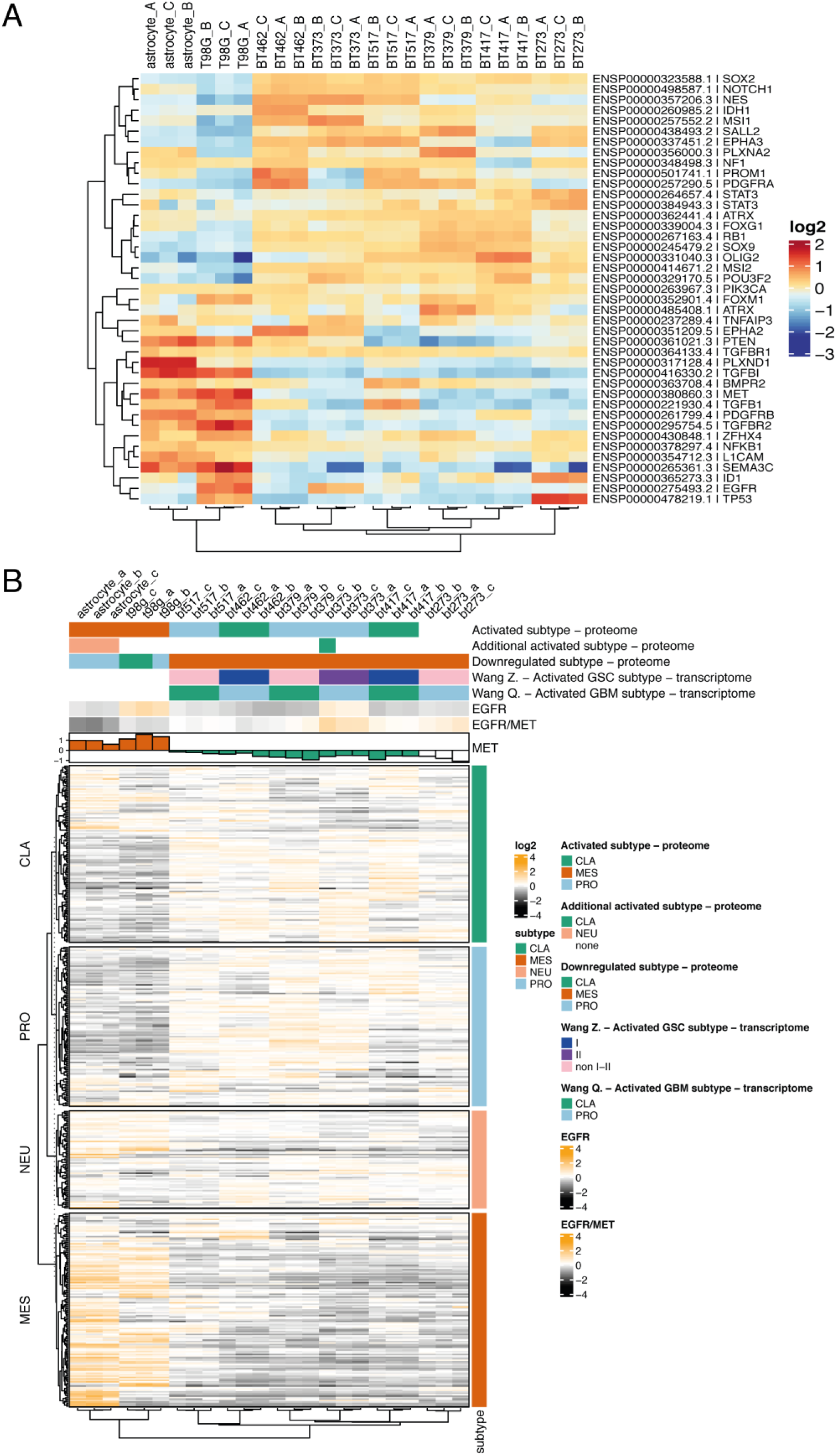
Protein expression of GSC markers described in literature. **A.** Hierarchical clustering (distance: 1-Spearman’s correlation coefficient) of GSCs and controls (astrocyte and T98G line) based on relative expression of known GSC markers; **B.** Protein expression of genes included in the Verhaak (2010) GBM subtypes’ gene sets identified in this study. Hierarchical clustering (distance: 1-Spearman’s correlation coefficient) of GSCs based on protein expression of the gene sets. Annotation map – the Wang Q. refers to the GBM mRNA subtypes classification of the GSCs based on mRNA expression. Wang Z. classification refers to the recently proposed GSC classification to type I and type II (see DeBacco *et. al*, 2021). Abbreviations: CLA = classical, PRO = proneural, MES = mesenchymal, NEU = neural.

**Figure S2.**
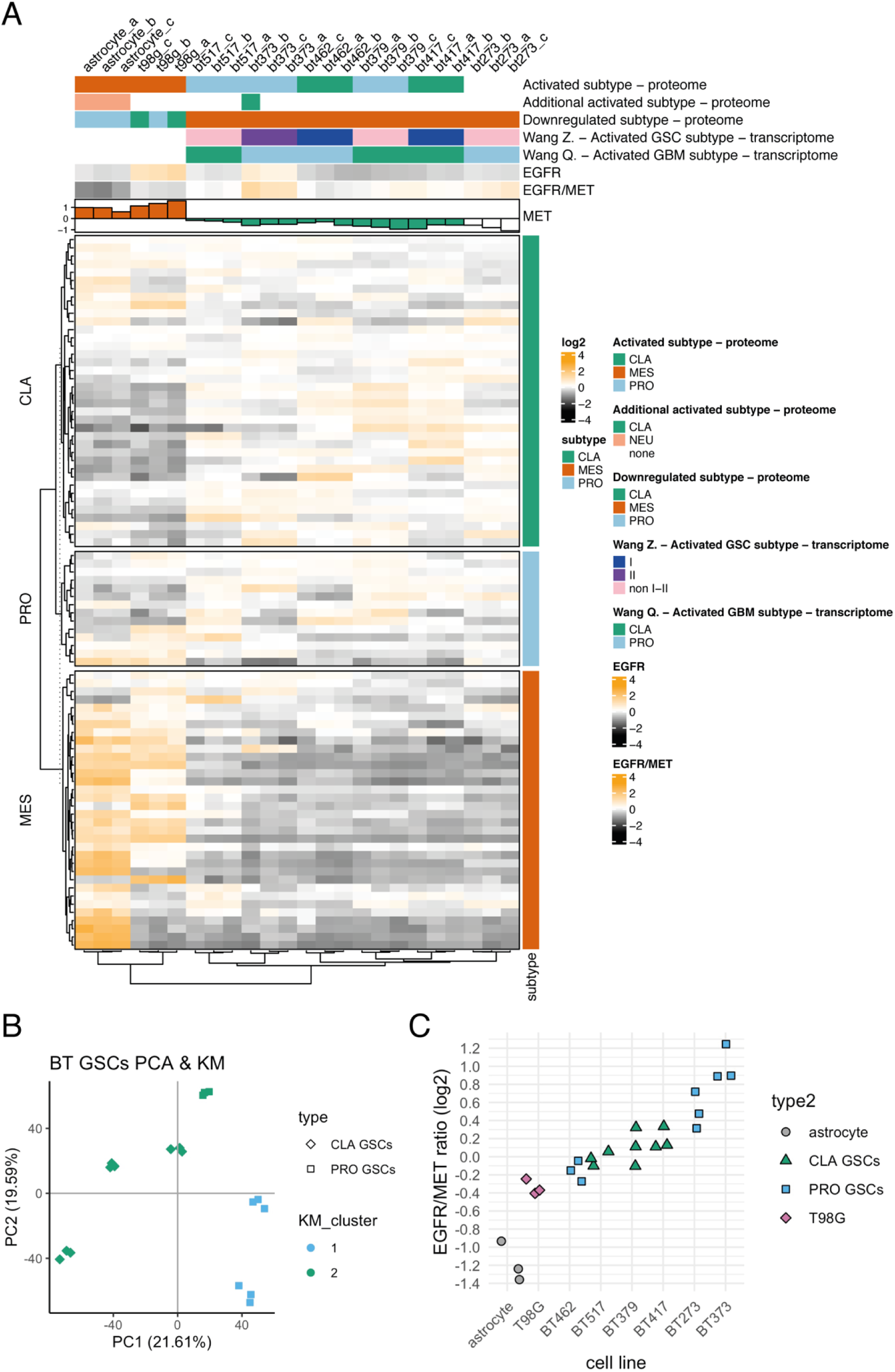
Protein expression of genes included in the Wang GBM subtypes’ gene sets. **A.** Hierarchical clustering (distance: 1-Spearman’s correlation coefficient) of GSCs and controls (astrocyte and T98G line) based on relative expression of the Wang GBM subtypes’ gene sets; **B.** For comparison - PCA and k-means (KM) clustering only of GSCs based on the expression of all the proteins without missing values. There was no clear separation between classical (CLA) and proneural (PRO) GSCs, as classified according to the Wang mRNA subtypes; **C.** EGFR/MET protein ratio on a log2 scale;

**Figure S3.**
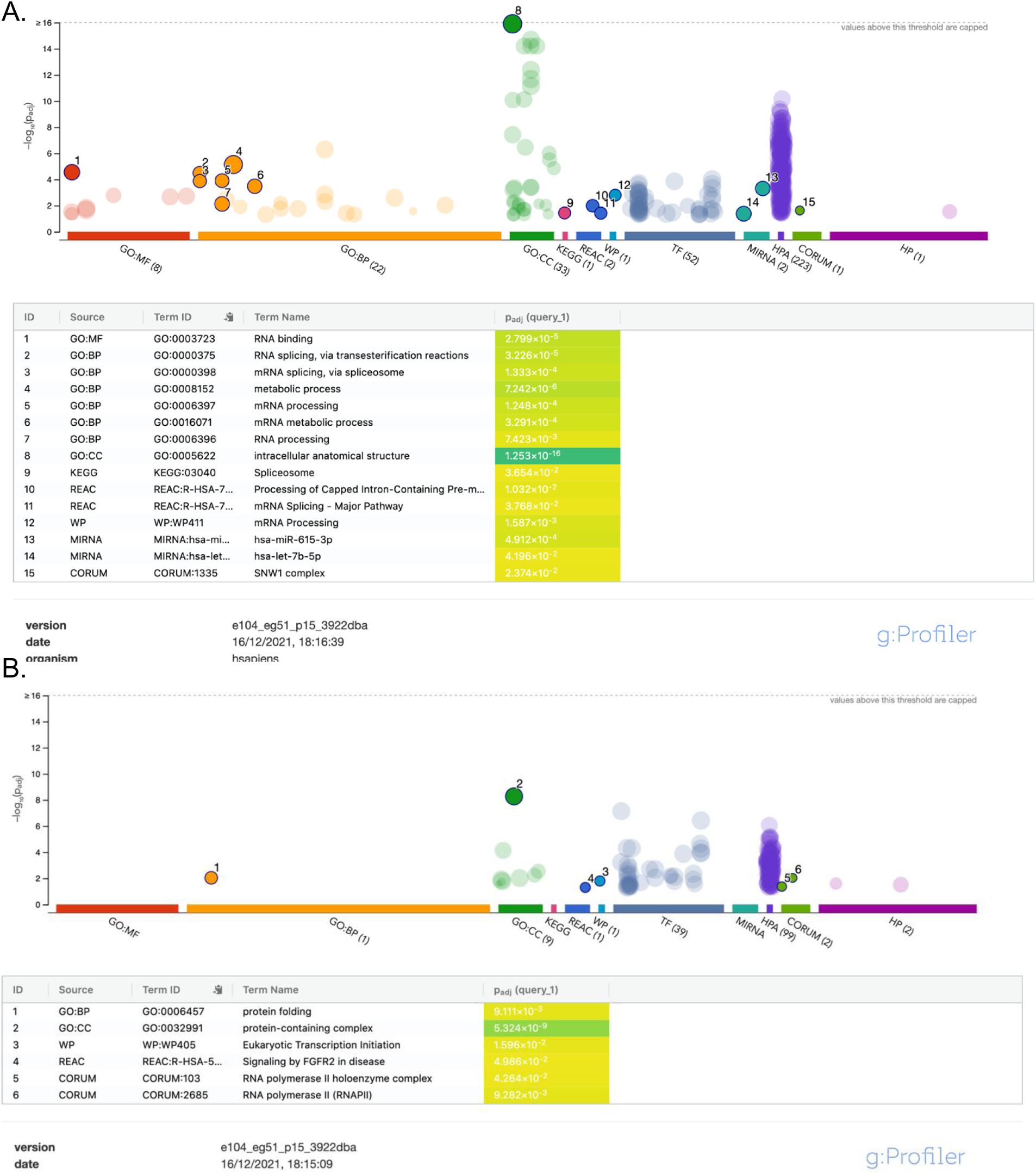
g:Profiler enrichment analysis of the genes that were outside of the 95% confidence intervals (CI) of the Bland-Altman plot comparing the agreement in mRNA-protein correlation estimates in GBM tissue and GSCs. **A.** Gene sets enriched in the list of proteins below the lower 95% CI, i.e. genes that had lower correlations in the GSCs compared to GBM tissue; **B.** Gene sets enriched in the list of proteins above the lower 95% CI, i.e. genes that had higher correlations in the GSCs compared to GBM tissue.

**Figure S4.**
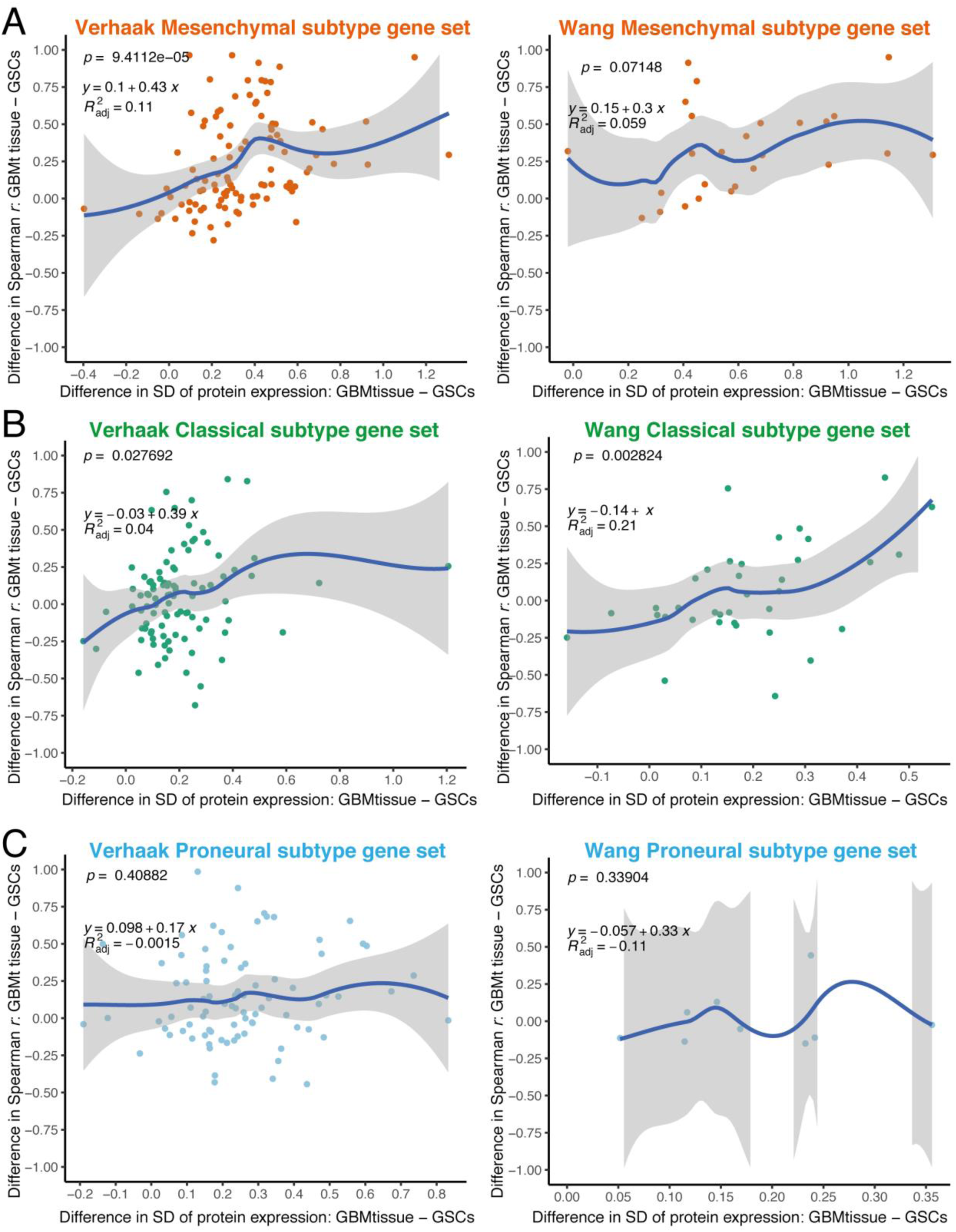
Per-gene mRNA-protein correlations in GSCs and GBM tissue. The difference between the standard deviations (SD) of protein expression in GBM tissue and GSCs (x axis) was associated with the difference in mRNA-protein correlations (Spearman r) estimated in GBM tissue and in GSCs (y axis). Higher proteins’ SD in GBM tissue was associated with higher correlation coefficients (r) in GBM tissue compared to GSCs for the mesenchymal **(A)** and classical **(B)** gene sets, but not for the proneural gene sets **(C)**.

**Figure S5.**
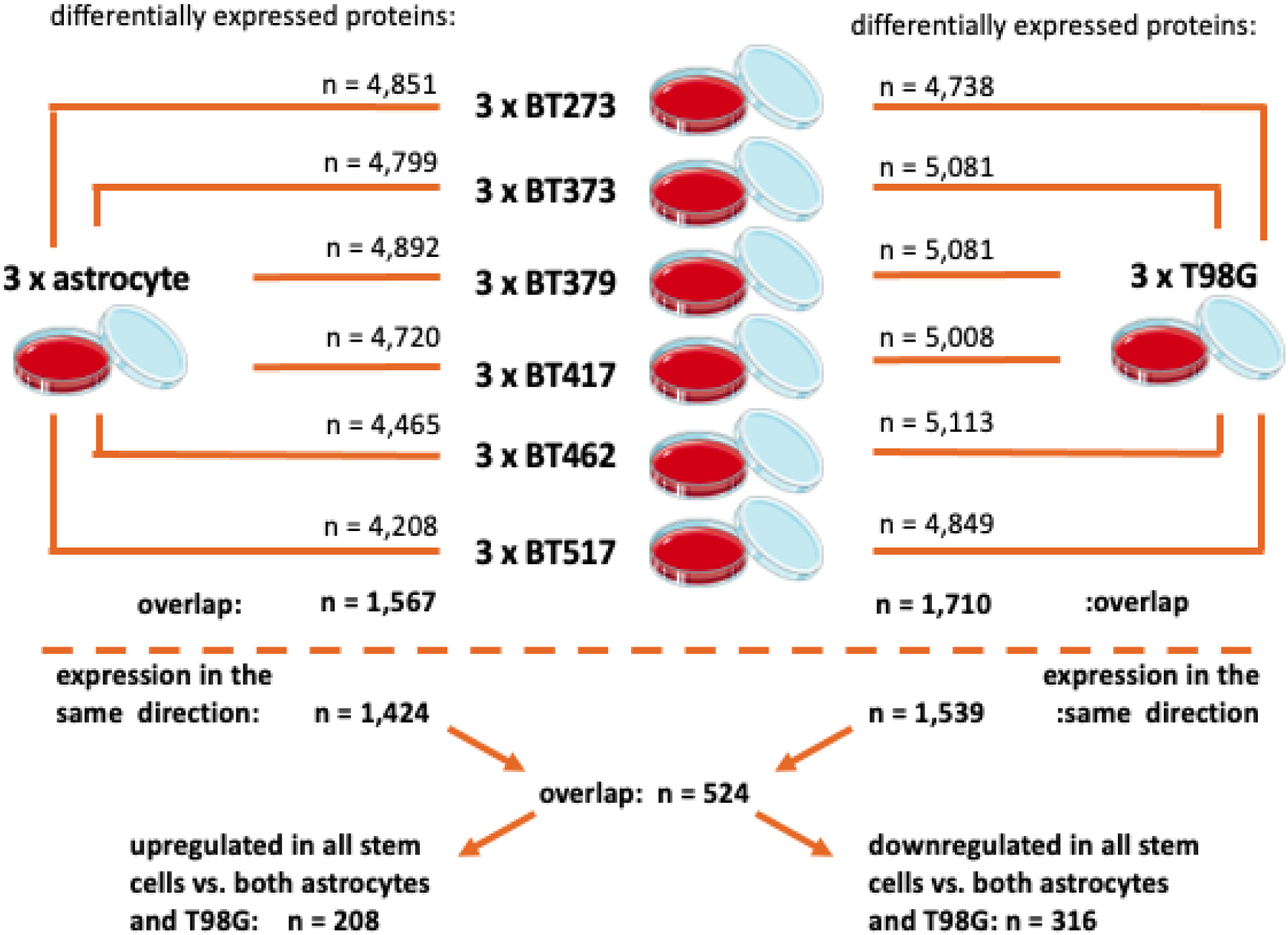
Differential expression algorithm for detecting GSAPS. Each GSC line was independently compared to the astrocyte and T98G line. Then we extracted the intersect of differentially expressed proteins in the same direction (over-/under-expressed in GSCs) in the comparison to the astrocyte and the T98G line, respectively. Finally, we took the intersect of consistently overexpressed and under-expressed proteins in GSCs that define GSAPS.

**Figure S6.**
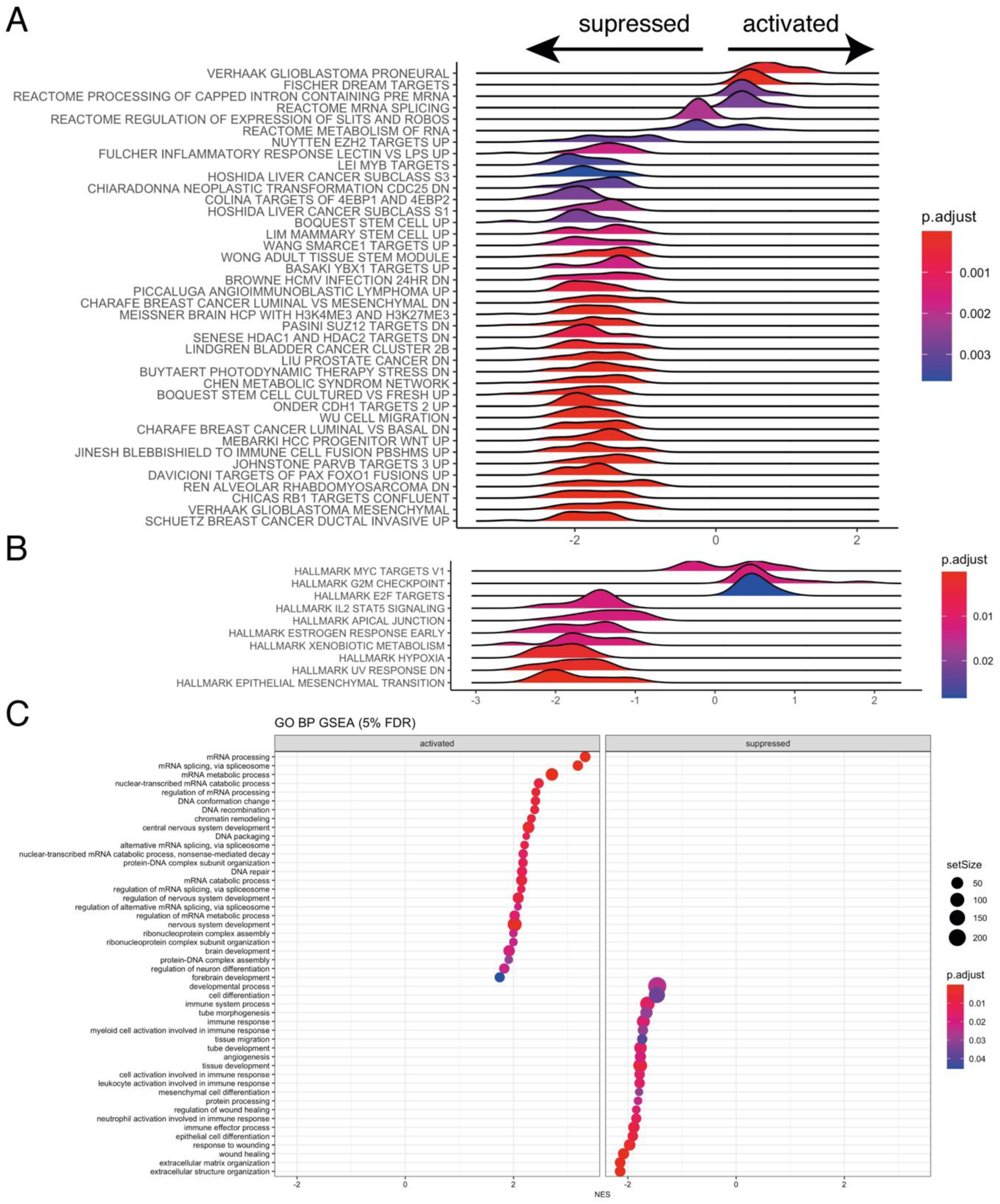
Gene set enrichment analysis (GSEA) of GSAPS, at 5% FDR. **A.** GSEA of chemical and genetic perturbations (GCP) and REACTOME gene sets included in the C2 collection of gene sets in the Molecular Signatures Database (MSigDB); **B.** GSEA of hallmark gene sets included in the H collection of gene sets in the MSigDB; **C.** GSEA of GO biological processes.

**Figure S7.**
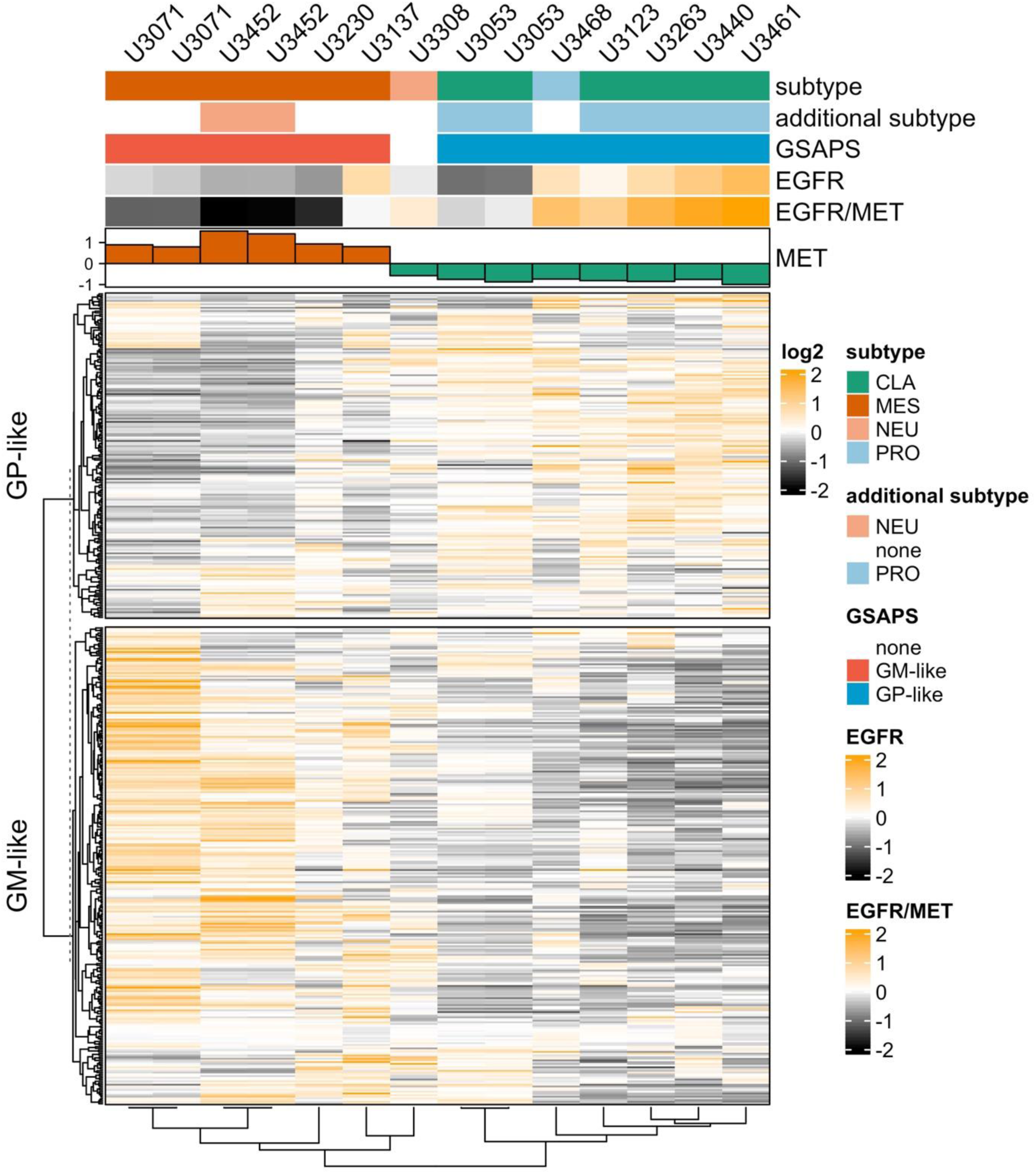
Hierarchical clustering of HGCC GSCs based on GSAPS protein expression. All classical GSCs had enrichment for the proneural subtype and the GPC-like GSAPS gene set, whereas the mesenchymal GSCs had enrichment for the GM-like gene set. Proneural + classical GSCs had a higher EGFR/MET ratio and lower MET levels, whereas the mesenchymal GSCs had a lower EGFR/MET ratio and higher MET levels.

**Figure S8.**
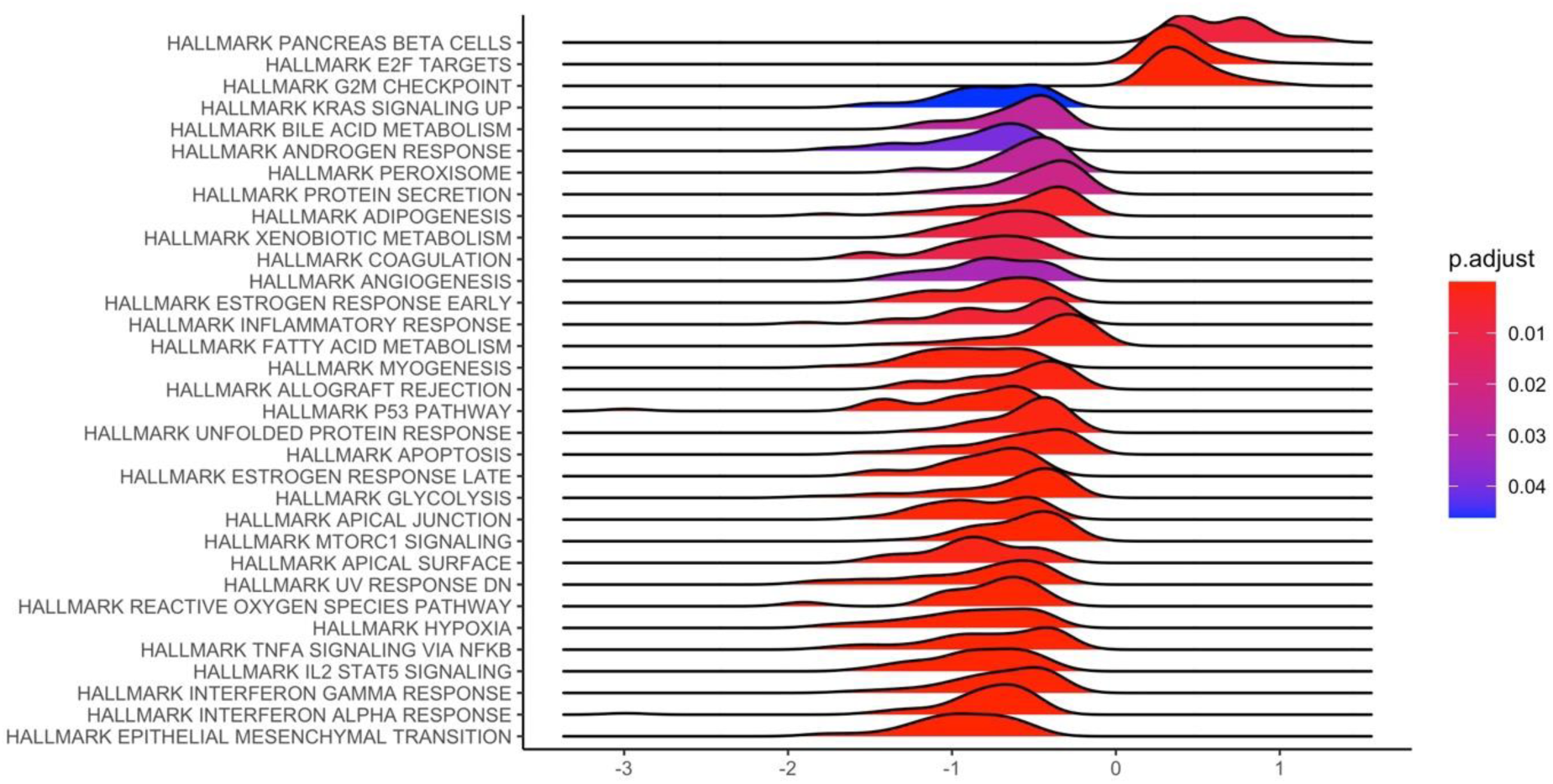
Gene set enrichment analysis (GSEA) of *hallmark* gene sets from the MSigDB, comparing protein expression of GPC-like GSCs to protein expression of GM-like GSCs, at 5% FDR.

**Figure S9.**
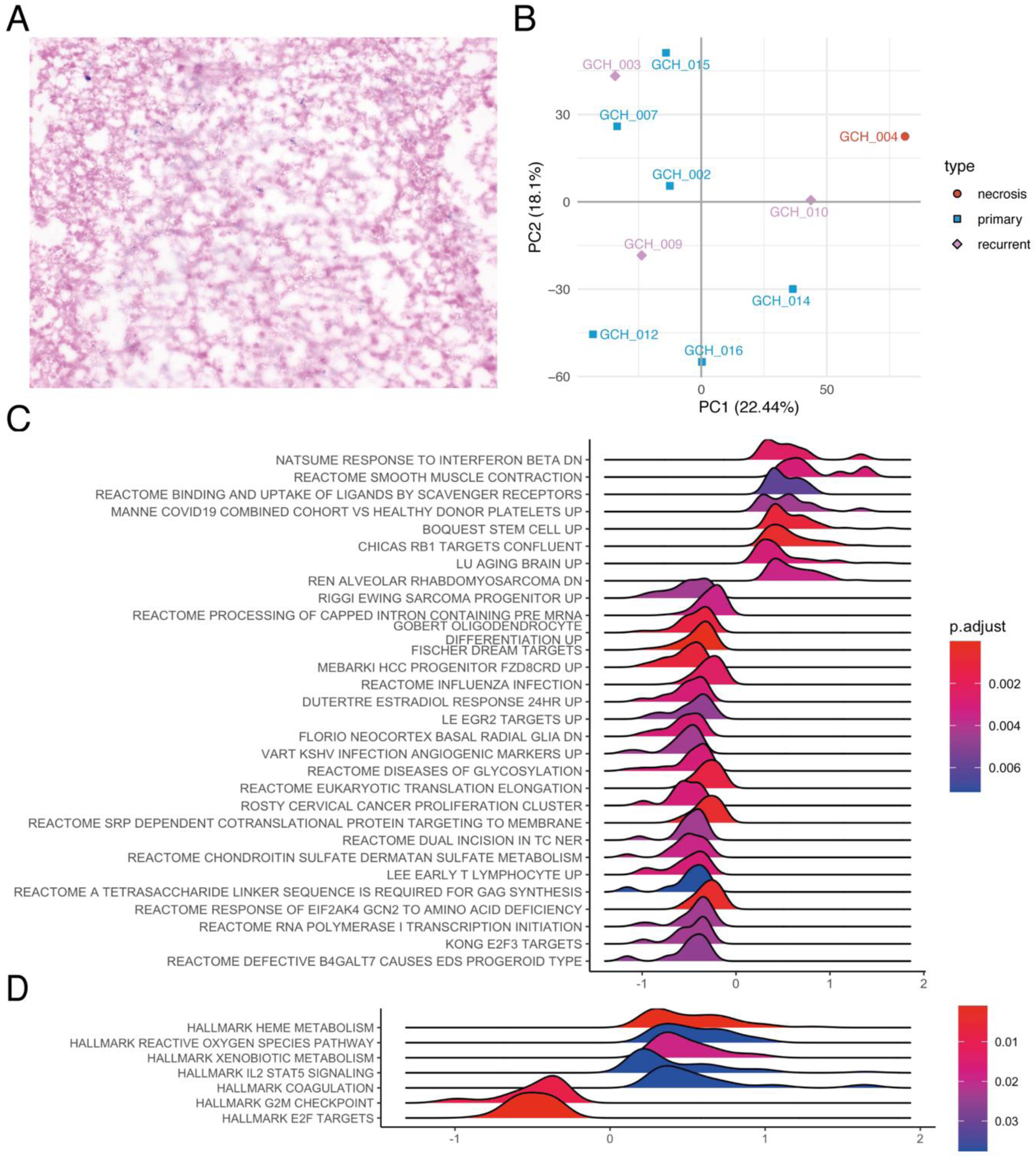
Pathways enriched in recurrent vs. primary GBM tumours. **A.** Haematoxylin and eosin staining of sample GCH004 showed extensive necrosis. The staining was repeated on another section; B. PCA clustering of GBM tissue samples based on bulk proteome expression; C. GSEA of C2 (subcategory: GCP and REACTOME) gene sets of the MSigDb, comparing recurrent to primary GBM tumours. The proteins were ranked based on a mean log2-FC comparing recurrent (n = 3) to primary (n = 6) GBM samples; D. GSEA of hallmark (H) gene sets of the MSigDb, comparing recurrent to primary GBM tumours.

**Figure S10.**
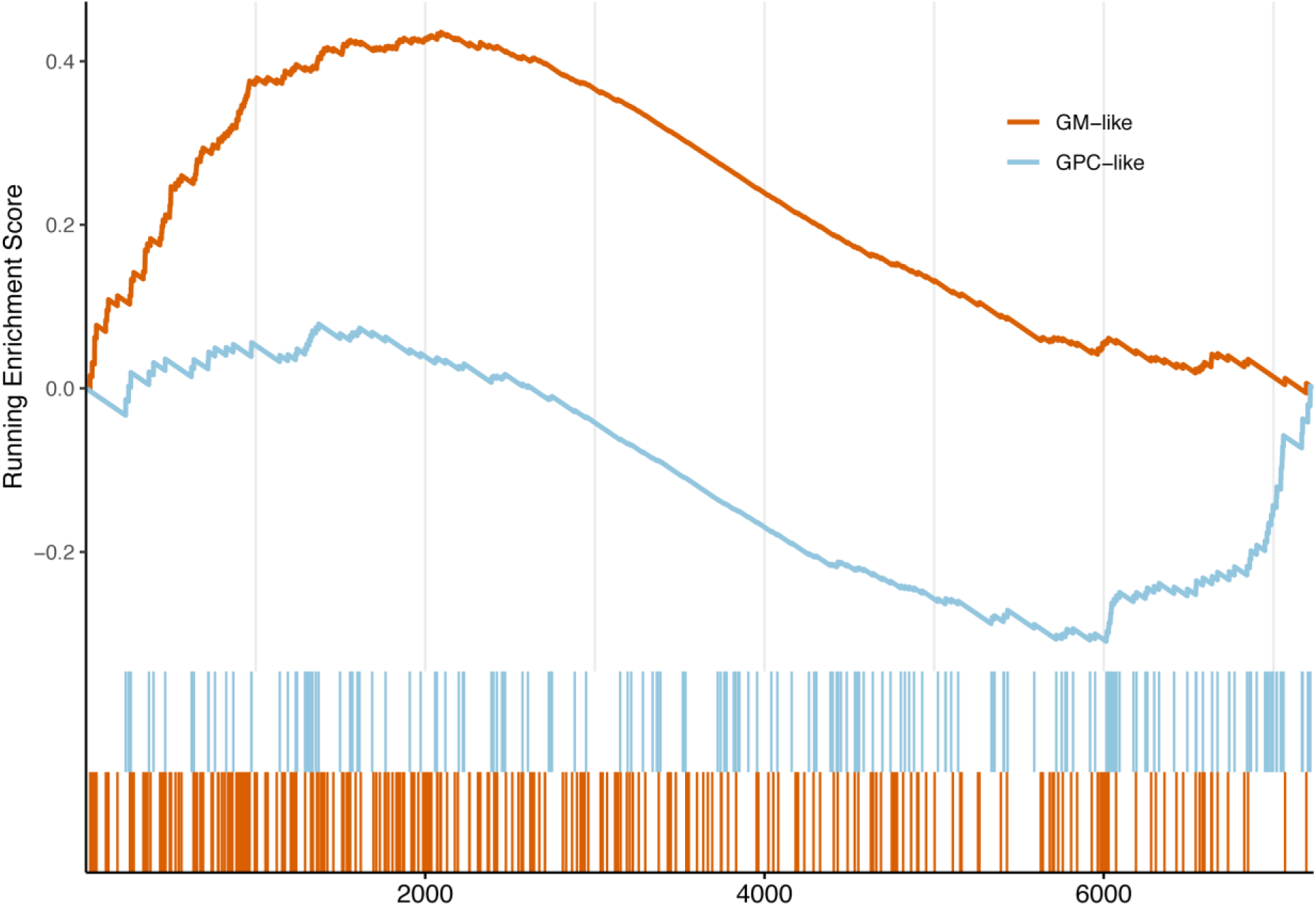
Single-sample GSEA of the GSAPS gene sets in the necrotic sample (p < 0.001, 1% FDR). Although the GM-like gene set was upregulated in the necrotic GBM tumour and the GP-like was supressed, the signals were not that consistent.

**Figure S11.**
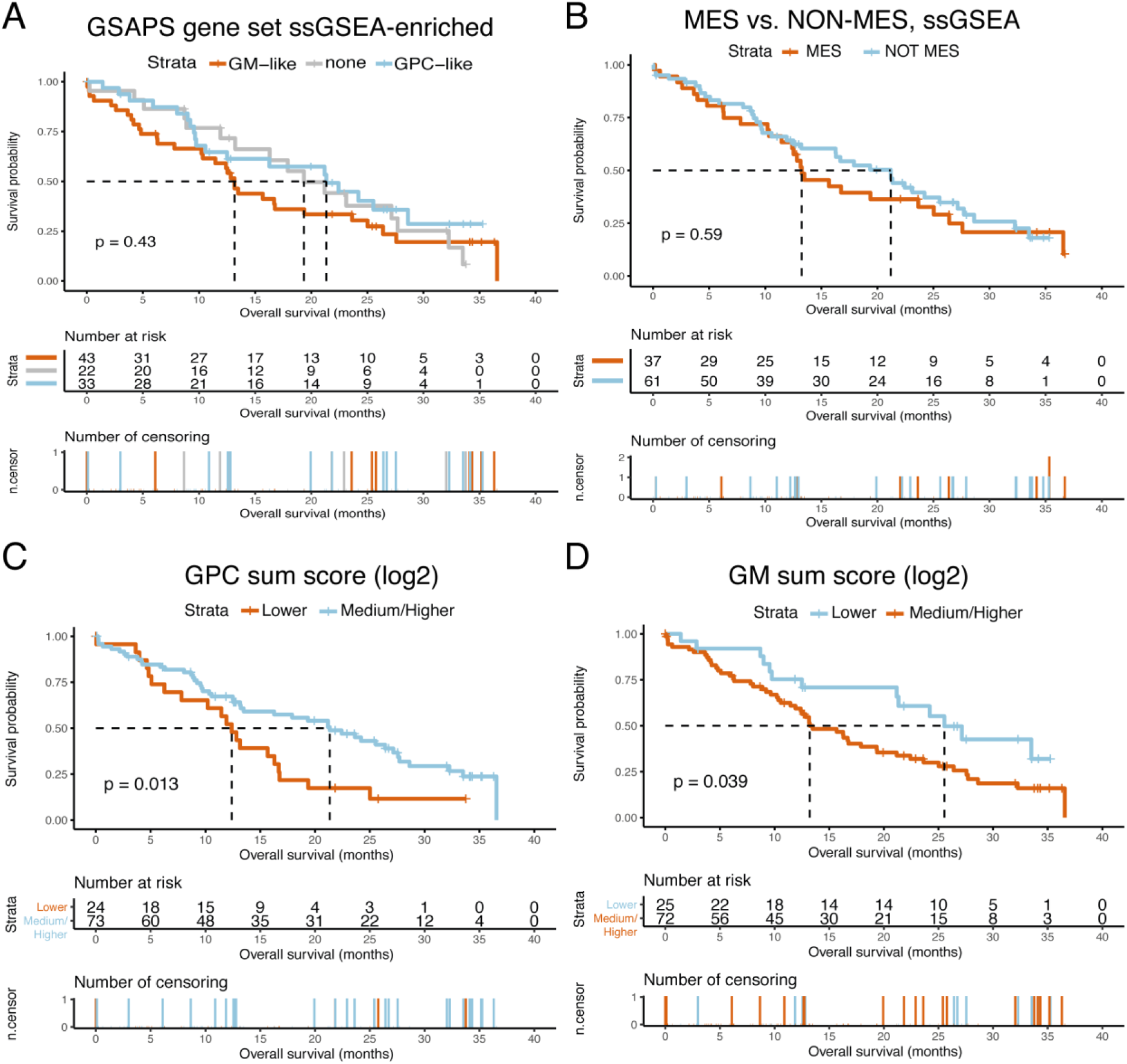
Overall survival in GBM patients based on GSAPS, Kaplan-Meier (KM) curves, CPTAC data. **A.** KM curves showing survival differences in patients categorised based on GSAPS gene set enrichment in GBM tissue; B. KM curves showing survival differences in patients with mesenchymal (MES) vs. non-mesenchymal (NON-MES) GBM (Wang gene sets); C. KM curves showing survival differences in patients categorised based on log2 GPC protein sum score expression to group of low (< first quartile) and medium/high (> first quartile) scores; D. KM curves showing survival differences in patients categorised based on log2 GM protein sum score expression to group of low (< first quartile) and medium/high (> first quartile) scores; The p values are based on logrank tests; the dashed lines present the median overall survival in the corresponding groups.

